# Interactomic affinity profiling by holdup assay: acetylation and distal residues impact the PDZome-binding specificity of PTEN phosphatase

**DOI:** 10.1101/2020.07.01.181487

**Authors:** Pau Jané, Gergő Gógl, Camille Kostmann, Goran Bich, Virginie Girault, Célia Caillet-Saguy, Pascal Eberling, Renaud Vincentelli, Nicolas Wolff, Gilles Travé, Yves Nominé

## Abstract

Protein domains often recognize short linear protein motifs composed of a core conserved consensus sequence surrounded by less critical, modulatory positions. Here we used an accurate experimental approach combining high-throughput holdup chromatographic assay and fluorescence polarization to measure quantitative binding affinity profiles of the PDZ domain-binding motif (PBM) of PTEN phosphatase towards the 266 known human PDZ domains. Inclusion of N-terminal flanking residues, acetylation or mutation of a lysine at a modulatory position significantly altered the PDZome-binding profile of the PTEN PBM. A specificity index is also introduced to quantify the specificity of a given PBM over the complete PDZome. Our results highlight the impact of modulatory residues and post-translational modifications on PBM interactomes and their specificity.

## Introduction

PDZs, named from the three proteins PSD-95, DlgA and ZO1, are globular protein domains that adopt a conserved antiparallel β-barrel fold comprising 5 to 6 β-strands and 1 to 2 α-helices. PDZ domains are involved in diverse cellular activities, such as cell junction regulation, cell polarity maintenance or cell survival [1]. PDZs recognize short linear motifs (called PDZ Binding Motif or PBMs) that follow particular sequence requirements and are mostly located at the extreme carboxy terminus of target proteins [2]. The human proteome contains 266 PDZ domains dispersed over 152 proteins [3] and thousands of presumably disordered C-termini matching a PBM consensus [4].

The core of a C-terminal PBM is formed by four residues, which are disordered in the unbound state but form, upon binding, an anti-parallel β-strand that inserts between a β-strand and a α-helix of the PDZ domain. A C-terminal PBM contains two conserved residues (positions are thereafter numbered backwards from the C-terminus, starting at p-0): a hydrophobic residue at p-0 and a characteristic residue at p-2, which actually determines the PBM classification: Ser / Thr for class I, a hydrophobic residue for class II and Asp / Glu for class III. Other positions located within or upstream of the core motif may also modulate the binding affinity ([5]–[8] and reviewed in [3]). In particular, systematic mutagenesis experiments have shown that amino acid replacements at positions −1, −3, −4 and −5, and sometimes even at upstream positions, can strongly alter the binding properties depending on the PDZ domain [9]–[11]. We and others have also shown that the length of the peptides or upstream or downstream sequences of the PDZ constructs used may influence the binding affinity in the assays [12]–[16].

Additionally, post translational modifications (PTM) at residues within or upstream of the PBM core are susceptible to alter the binding affinity for PDZ [17], and therefore the PDZ / PBM network. Protein acetylation is an example of PTM that mainly targets lysine residues. Acetyltransferases catalyze the transfer of an acetyl group from acetyl-coenzyme A to the ∊-amino group of a lysine residue, inducing the neutralization of the positive charge of the lysine side chain. The reaction can be reversed by lysine deacetylases. By modifying the chemical nature of the protein, the acetylation process may alter its binding properties. In particular, an acetylated protein may become “readable” by specialized acetyl-lysine binding domains such as bromodomains [18]. Acetylation occurs in a large variety of protein substrates and plays important roles in protein regulation, DNA recognition, protein / protein interaction and protein stability [19]. Originally widely described for histone proteins, it has also been observed for a growing number of non-histone proteins [20], such as PTEN [21].

PTEN is a lipid phosphatase protein located in the cell nucleus with a prominent tumor suppressor activity. When brought to the plasma membrane, PTEN is able to antagonize the phosphatidylinositol 3-kinase (PI3K), inhibiting the PI3K-dependent cell growth, survival and proliferation signaling pathways [22]. Interestingly, PTEN harbors a class I PBM –ITKV_COOH_– that appears to be critical for regulating its functions [23],[24]. The PDZ binding to the PTEN PBM leads to a stabilization of PTEN and an increase of its catalytic activity [25]. The PBM of PTEN presents several original characteristics. On the one hand, a structural study revealed an unconventional mode of binding of PTEN to the PDZ domain of the human kinase MAST2 [26]: while the core of the PTEN PBM displays a canonical interaction with the PDZ domain, a Phe residue at p-11 (F392) distal from the core PBM establishes additional contacts with MAST2 through a hydrophobic exosite outlined by β2- and β3-strands of the PDZ domain. On the other hand, lysine K402, located at the p-1 position of the PBM core in PTEN, has been suggested to represent a putative target of an acetylation reaction that might modulate PTEN binding to PDZ domains and thereby affects other PTEN activities [21]. Remarkably, those original characteristics of the PBM of PTEN (unconventional PDZ binding mode of PTEN and potential modulation by acetylation) have been examined only in context of interaction with a few PDZ domains. It is thereafter interesting to cover their impact on the interactome with the full PDZome, thus requiring the use of a high-throughput screening method, as the holdup.

The holdup method is a chromatographic approach in solution developed in our group that allows to measure the binding strength of a peptide, attached to a resin, against a library of domains of a same family. We initially proposed this method to explore the interaction between PBM peptides and the human PDZ domains [27]. Briefly, a soluble cell lysate containing individually overexpressed PDZ domain is incubated until equilibrium with a calibrated amount of streptavidin-resin saturated either with the target biotinylated PBM peptide or with biotin as a reference. The flow-throughs containing the unbound protein fraction are recovered by filtration and loaded on a capillary electrophoresis instrument to quantify the amount of remaining free PDZ. The stronger the steady-state depletion of the PDZ domain in the flow-through as compared to the reference, the stronger the PDZ / PBM binding interaction. The assay is particularly suited to quantitatively evaluate and compare large numbers of interactions. This method delivers, for each PBM / PDZ pair, a “binding intensity” (BI), whose value can in principle range from 0.00 (no binding event detected) to 1.00 (maximal binding event). The approach has been automated [28] and the human PDZ library was recently extended to the complete 266 PDZ domains known in human proteome [29]. The high-throughput assay is implemented on 384 well-plates, and can probe a single peptide in triplicate or up to 3 different peptides in singlicate against the 266 PDZ domains. The full processing leads to a binding profile, i.e. a list of binding strengths in decreasing order exhibited by a given PBM towards the entire PDZome. The high accuracy and efficiency of the holdup assay has been validated previously [4],[15],[28],[30]. Very recently, a manual version of the holdup assay with purified samples and using widespread benchtop equipment has been implemented and has proven to be reliable [31].

In the present work, we investigated how the acetylation at position K402 in PTEN (−IT^Ac^KV_COOH_− thereafter corresponding to p-1 position in the PBM), would alter the binding affinity profile of the PTEN C-terminus to the full complement of known human PDZ domains (the PDZome). We also assessed the contribution of the K402R mutation, expected to preserve the positive charge and the overall bulkiness of the lysine residue, as well as the effect of the presence of the hydrophobic residue at p-11 (F392). For these purposes, we combined the updated high-throughput holdup assay with fluorescence polarization (FP) measurements allowing to convert each BI value into affinity. We obtained all the affinities of the complete human PDZ library for wild-type, acetylated and mutated versions of the PBM of an 11-mer PTEN C-terminal peptide as well as an extended 13-mer peptide. We also introduced a tentative “promiscuity index” to quantify the PDZome-binding specificity of each peptide. The results show that acetylation affects the affinities for the PDZome and highlight the importance of the exosite in modulating the PDZome specificity for the PDZ-binding motif of PTEN.

## Material and Methods

### Protein expression and purification

The 266 PDZ domains that constitute the used PDZ library (“PDZome V.2”) were produced using constructs with optimized boundaries as described previously [32]. All the genes were cloned into pETG41A or pETG20A plasmid. The expressions in *E.coli* resulted in a recombinant protein fused to an N-terminal solubility tag (His-MBP or TRX). The expressed tag-PDZ concentrations were quantified using capillary gel electrophoresis and cell lysates were diluted to reach approximately 4 μM tag-PDZ before freezing in 96-well plates. A detailed protocol of the PDZ library production, expression and benchmarking can be found in [29]. PDZ domains are named according to their originating protein name followed by the PDZ number (e.g. NHERF1-1 as the first PDZ domain of the NHERF1 protein).

For FP assay, tandem affinity purified His_6_-MBP-PDZ proteins were used. Cell lysates were purified on Ni-IDA columns, followed by an MBP-affinity purification step. Protein concentrations were determined by far-UV absorption spectroscopy. A detailed protocol has been published previsouly [4].

### Peptide synthesis

All 11-mer biotinylated peptides (PTEN_11: DEDQHTQITKV, PTEN_Ac: DEDQHTQIT***ac*K**V and PTEN_KR: DEDQHTQIT**R**V) were chemically synthesized on an ABI 443A synthesizer with Fmoc strategy by the Chemical Peptide Synthesis Service of the IGBMC, while PTEN_13 (**PF**DEDQHTQITKV) was purchased from JPT Innovative Peptide Solutions with 70%–80% purity. A biotin group was systematically attached to the N-terminal extremity of the peptide via a TTDS linker while fluorescent peptides were prepared by directly coupling fluorescein to the N-terminus. Predicted peptide masses were confirmed by mass spectrometry. Due to the lack of aromatic residue, peptide concentrations were first estimated based on the dry mass of the peptide powders and subsequently confirmed by far-UV absorption (at 205 and 214 nm).

### Holdup assay

The Holdup assay was performed in singlicate for the three 11-mer PTEN variants and the 13-mer PTEN variant as described in [28],[29]. Prior to interaction assay, the streptavidin resin was saturated with biotinylated PBM peptides and then washed with an excess of free biotin, while the reference resin was incubated only with biotin. Right before the holdup experiment, the PDZ library was spiked with an internal standard of lysozyme. Then, the biotin- or PBM-saturated resins were incubated with diluted cell lysates, each in a distinct well of a 384-well plate, allowing to adjust the concentration of tag-PDZ at around 4 μM. After a sufficient time for the complex to form (15 min.), a fast and mild filtration step is performed and the tag-PDZ concentrations were measured by capillary electrophoresis instrument (LabChip GXII, PerkinElmer, Massachusets, USA). A detailed protocol of how to run the holdup assay in an automatic way using liquid handling robots can be found in [29]. Standard markers were used to convert migration time into molecular weight on the LabChip software and inappropriate molecular weight markers were corrected or excluded.

### Holdup data quality check and processing

Holdup data can be missing for some tested pairs mainly for three reasons: i/ biochemical issues, specially when the over-expressed domain is not concentrated enough in the sample, ii/ acquisition problems mainly because of a misreading of the Caliper data, iii/ technical difficulties related to data processing. For points i/ and ii/, many efforts have been made to optimize the expression and to run the LabChip GXII instrument in the best conditions. For point iii/, we developed bioinformatics processing tools in order to improve the accuracy and reproducibility of the intensity measurement of the tag-PDZ peak in the chromatogram [33]. Briefly a baseline correction of the electropherograms is first performed in order to remove the background noise and extract the real intensities using Python package available in https://spikedoc.bitbucket.io under the name of SPIKE.py [34],[35]. Then intensities are normalized using the internal standard (lysozyme as previously mentioned) to correct potential variations over all the protein concentrations. Lastly, both the sample and the reference electropherograms were superimposed by adjusting the molecular weight on the X-axis according to a linear transformation (translation and dilation) of the sample electropherogram as compared to the reference one.

Beyond the purpose of this article, we have accumulated several tens of thousands of PDZ / PBM interaction data with the holdup protocol used here. Experienced holdup data curators combined four quantitatively evaluable quality criteria to retain or discard data during visual inspection. Individual electropherograms must display a sufficiently high intensity of the normalization peak (criterium 1) and of the tag-PDZ peak (criterium 2) while the signal of crude extract should be kept as low as possible compared to the tag-PDZ peak (criterium 3) (**Fig. 1A**). When comparing two electropherograms, the elution profiles must be sufficiently aligned (criterium 4) (**Fig. 1B**). In order to rationalize and accelerate data curation, we assigned to each criterium an individual quality score ranging from 0 to 1 from the lowest to the highest quality data (**Fig. 1C**). To avoid a cut-off effect, a linear or quadratic transition was introduced depending on the quality criteria type. The product of the resulting individual scores led to a global quality score in the 0-to-1 range. We calculated such scores for holdup data sets that had been treated by expert curators, then compared the scores of the data that had been either rejected or retained by the curators. This allowed us to semi-empirically set a threshold value of 0.6 which maximizes the true positive rate and minimizes the false negative rate. This threshold was automatically used to distinguish data to be rejected from those to be retained in a way that generally agrees with the expert curator’s decision. For the datasets used in the present study, the percentage of rejection never exceeded 10%.

**Fig 1.**
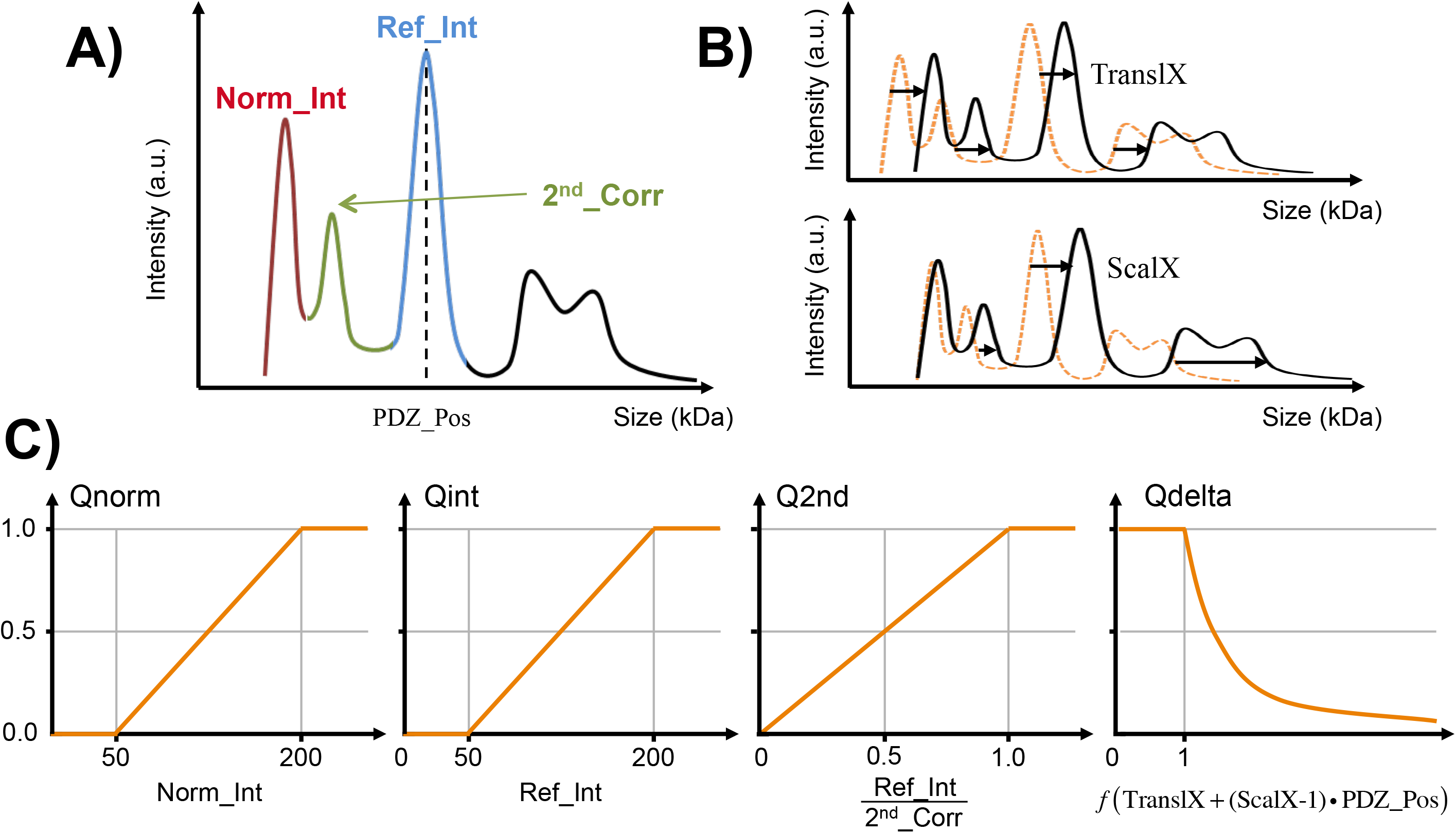
Quality criteria and their conversion to the individual quality scores used to filter the holdup data. (**A**) A schematized electropherogram showing intensities of the normalization peak (*Norm_Int*) and of the MBP-PDZ peak (*Ref_Int*) visible in the red and blue regions, respectively. The region in green corresponds to the proteins of the crude extract, which is supposed to be kept low as compared to *Norm_Int* and *Ref_Int* in order to ensure that the MBP-PDZ is not underexpressed **(B)** The linear transformation used to superimpose the sample and reference electropherograms should be as neutral as possible: the *TranslX* translation factor and the *ScalX* scaling coefficient (>1 for dilation or <1 for a contraction) should be as close as possible to 0.0 and 1.0, respectively. **(C)** Profiles of the individual quality scores used to filter the data. In order to ensure that the analyzed samples were not too diluted, the scores vary linearly between 0 (low quality) and 1 (high quality) for the intensity of the normalization peak (Q_norm_) or the MBP-PDZ peak (Q_int_). Q_2nd_ is a quality score allowing to reject samples with low MBP-PDZ expression while Q_delta_ combines the *TranslX* and *ScalX* parameters and varies exponentially. *Double column fitting image*.

For filtered data, the BI was extracted with the following equation (**Eq. 1**) that estimates the depleted fraction after superimposition of the sample and reference electropherograms:

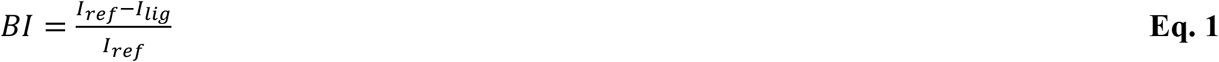

where I_ref_ and I_lig_ are the intensities of the tag-PDZ peaks measured in the reference and the sample electropherograms, respectively, for a given PDZ domain / PBM peptide interaction pair.

Data reproducibility has been previously explored for several PDZ / PBM pairs resulting in a standard error of the mean of about 0.07 BI unit (data not shown + [28]). This suggests that the maximal BI values differ significantly from PTEN_Ac or PTEN_KR constructs as compared to PTEN_11 and in a less extend to PTEN_13. In some cases, negative BI values as low as −0.20 can be observed and seem to be reproducible (data not shown). This could result from a lower intensity of the reference PDZ / PBM peak as compared to the sample PDZ / PBM peak, potentially due a preference of the PDZ domain for beads fully saturated with biotin as compared to beads with biotinylated peptide. As reported previously, we have also investigated the limit of detection by repeating the holdup experiments for an irrelevant “neutral” peptide owing no specific PBM consensus sequence. Almost all BI values were below 0.10 (98% of all measured PDZ / PBM pairs) and showed a standard deviation of less than 0.10 (considering 95% of the data) [28]. According to this, we applied a conservative safety factor of 2 that leads to a limit for BI of 0.20. This cut-off represents a very stringent threshold retaining only high-confidence PDZ / peptide interactions, and eliminating most of the false positives.

### Steady-state fluorescence polarization

FP data were measured in 384-well plates (Greiner, Frickenhausen, Germany) using a PHERAstarPlus multi-mode reader (BMG labtech, Offenburg, Germany) with 485 ± 20 nm and 528 ± 20 nm band-pass filters for excitation and emission, respectively. N-terminal fluorescein-labeled HPV16E6 (fluorescein-RRETQL), RSK1 (fluorescein-KLPSTTL) and phospho-RSK1 (fluorescein-KLPpSTTL) were used as tracers. In competitive measurements, the 50 nM fluorescent reporter peptide was first mixed in 20 mM HEPES pH 7.5 buffer (containing 150 mM NaCl, 0.5 mM TCEP, 0,01% Tween 20) with the PDZ domain at a sufficient concentration to achieve high degree of complex formation. Subsequently, increasing amount of unlabeled peptide was added to the reaction mixture with a total of 8 different peptide concentrations (including the 0 nM peptide concentration i.e. the absence of peptide). Titration experiments were carried out in triplicate. The average FP signal was used for fitting the data to a competitive binding equation with ProFit, an in-house Python-based program [36], allowing to extract the apparent affinity values. In our competitive assays, every tested PDZ domain detectably bound to at least one PBM peptide, in agreement with well folded PDZ domains.

### Conversion from BI values to dissociation equilibrium constants

BIs were transformed into dissociation constants (K_D_) using the following formula:

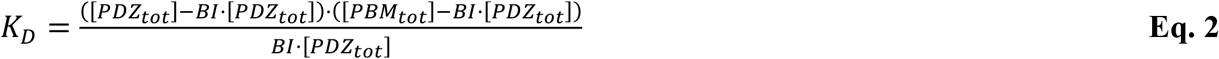

where [PDZ_tot_] and [PBM_tot_] correspond to the total concentrations of the PDZ domain (usually around 4 μM) and the PBM peptide used during the assay. Since the PBM_tot_ concentration in the resin during the holdup assay parameter may differ from one peptide to another and remains unknown, it is impossible to directly convert BI values into K_D_ constants. To extract the PBM concentration, we systematically determined by FP the K_D_ constants for a subset of PDZ / PBM pairs that were used to back-calculate the peptide concentrations in the holdup assays when quantifiable and significant (>0.20) BI values were available for the same pairs (**Eq. 2**). For each PBM, the average peptide concentration was calculated after outlier rejection based on the absolute distances from the median as compared to three times the standard deviation (3σ rule), with never more than 2 values rejected.

## Results

### An experimental strategy to measure large numbers of reliable affinity data

For this study, we wished to generate accurate and complete PDZome-binding affinity profiles for four peptide variants of the C-terminal PBM of PTEN. In practise, this requires measuring the individual affinities of 4×266=1064 distinct PBM-PDZ pairs. A singlicate holdup experiment is well suited for such a task. Taking into account the additional ~360 biotin-PDZ negative control measurements required for data treatment, the assay delivers ~1400 filtrates of protein extracts, which must each be individually subjected to capillary electrophoresis. Next, individual electropherograms must be visually curated and analyzed by an expert user to extract the binding intensities (BI) values that will compose the final profiles. As described in the material and methods section, we rationalized the data curation step by introducing a numerical global quality score. Since the assay requires expensive materials and labor-intensive data treatment, one should favor an approach based on singlicate holdup runs. We therefore used a strategy that combines one holdup assay run in singlicate with a medium-throughput competitive FP protocol run on a large proportion of the PDZ / PBM interacting pairs detected in the holdup assay (see material and methods). This strategy warrants the obtention of highly reliable affinity data for all PDZ / PBM interacting pairs that pass the quality score filtering step after the holdup assay. Representative holdup data recorded for one PBM (PTEN_11) are shown in **Fig. 2A**. After normalization of the two capillary electropherograms recorded for both the PBM of interest and the biotin reference, the comparison of the intensities of the two resulting PDZ peaks informs about the strength of the interaction: the stronger the depletion, the stronger the binding. Representative FP data are shown in **Fig. 2B**. The apparent affinities were obtained by fitting the anisotropy data considering a competitive binding model [37]. The holdup BI values and the binding strength derived from competitive FP measurements are consistent: higher the BI, stronger the affinity.

**Fig 2.**
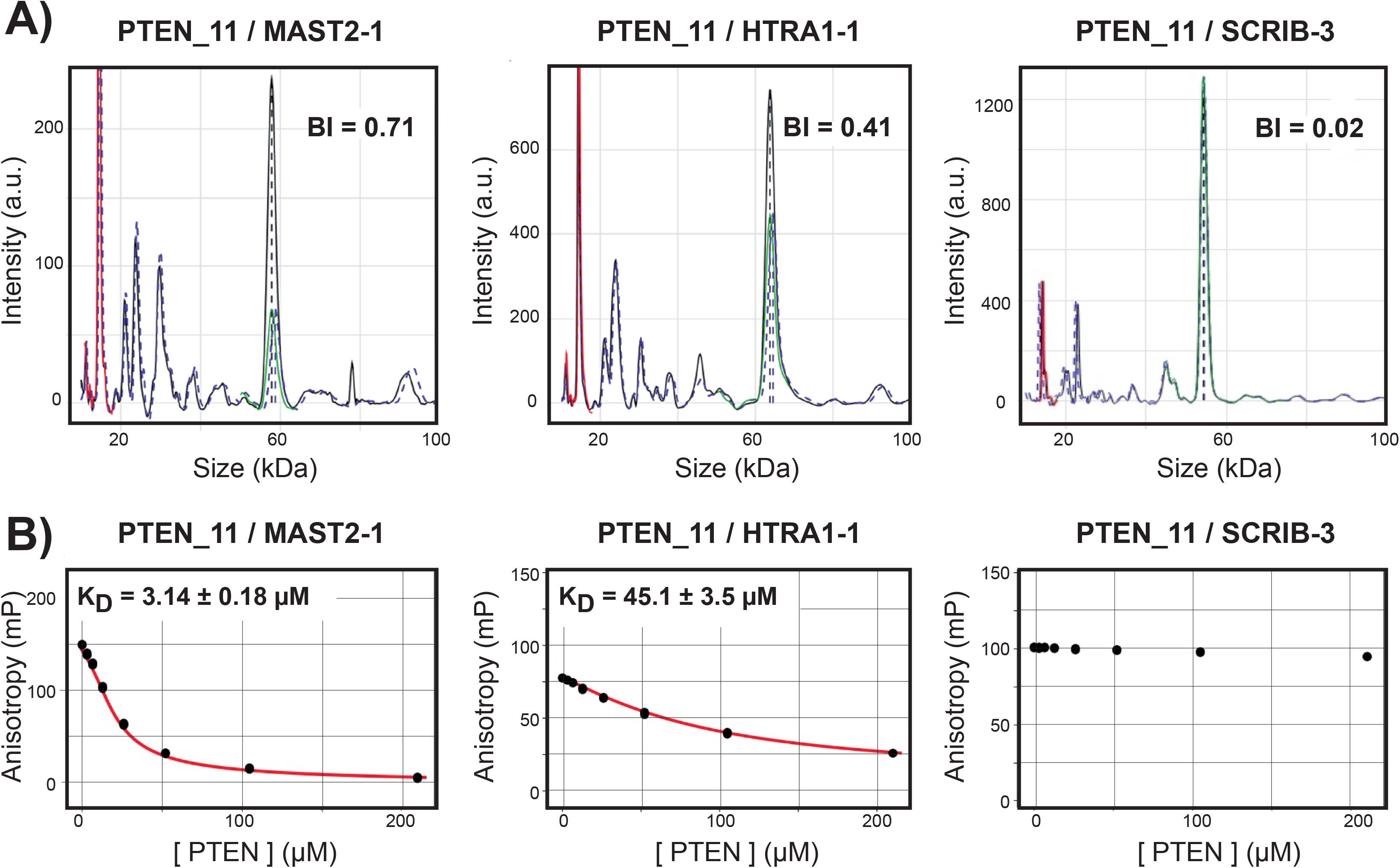
Complementarity of holdup and fluorescent polarization data. The interaction data of PTEN_11 with MAST2-1, HTRA1-1 and SCRIB-3 are shown as examples of strong affinity, weak affinity or non-binding, respectively, all measured by holdup (**A**) and FP (**B**) methods. (**A**) After superimposition of the two electropherograms recorded for the PBM of interest (blue dotted line) and for the biotin reference (black solid line), the normalization of the electropherogram of the PBM compared to the one of the reference is done using the signal of the lysozyme added in every sample at a constant concentration (red peak). The region between 20 and 60 kDa which contains peaks of the crude extract supposedly to be constant, is used to verify the proper intensity normalization of the two electropherograms. The intensities of the peak of interest after proper alignment along the molecular weight scale (region covered by the green dotted line) are subsequently used to quantify the depletion of an individual PDZ domain. All those normalization and alignment steps are performed automatically and are important as the electropherogram overlap is never perfect. The holdup ultimately delivers “binding intensities” (BI) for each PBM/PDZ interaction pair, which in principle vary in a range from 0.00 (no binding) to 1.00 (strong binding). (**B**) In competitive FP measurements, polarization signal was recorded for increasing amounts of unlabeled peptide added to a solution of pre-formed PDZ / labeled peptide complex. The complexes consisted of MAST2-1, HTRA1-1 and SCRIB-3 mixed with 50 nM of labeled fpRSK1, fRSK1 and f16E6 peptides, respectively. The PDZ concentration depends on each sample and is adjusted to reach >50-80% complex formation to ensure a satisfactory signal-to-noise ratio. Each panel shows the average of three titration curves (black dots) and the fit results (red curves with the apparent K_D_ values) using competitive binding model. *Single column fitting image.*

### Generating PDZome-binding BI profiles of the four PTEN variant PBMs by holdup assay

We applied the holdup assay to generate PDZome-binding profiles of three 11-mer peptides (PTEN_11 for the native sequence, PTEN_Ac and PTEN_KR for the acetylated and K402R mutated version of PTEN_11, respectively), as well as an extended 13-mer peptide (PTEN_13). Considering the quality score filtering step, we managed to quantify the interactions of 213, 233, 215 and 257 PDZ for the PTEN_11, PTEN_Ac, PTEN_KR and PTEN_13 peptides, respectively, which corresponds to 80%, 81%, 88% and 97% of the human PDZome. All holdup plots that detected a binding event with a binding intensity BI>0.20 are shown in **Supp. Fig. S1**. The four resulting holdup datasets were then plotted independently in the form of “PDZome-binding profiles” representing the individual BI values versus the PDZ domains ranked from higher to lower BI values (**Fig. 3**). PTEN_11 showed a maximal BI value of 0.71, i.e. a lower binding strength as compared to the ones of PTEN_KR, PTEN_Ac or PTEN_13 (BI = 0.86, 0.90 and 0.81, respectively). Using BI>0.2 as a minimal threshold value for retaining high-confidence PDZ / peptide interactions, the holdup assay identified 19, 43, 37 and 24 PDZ domains as potential binders for the PTEN_11, PTEN_Ac, PTEN_KR and PTEN_13 peptides, respectively. Altogether, they represent a total of 123 potential binders, of which 60 are non-redundant PDZ domains distributed over 46 distinct proteins.

**Fig 3.**
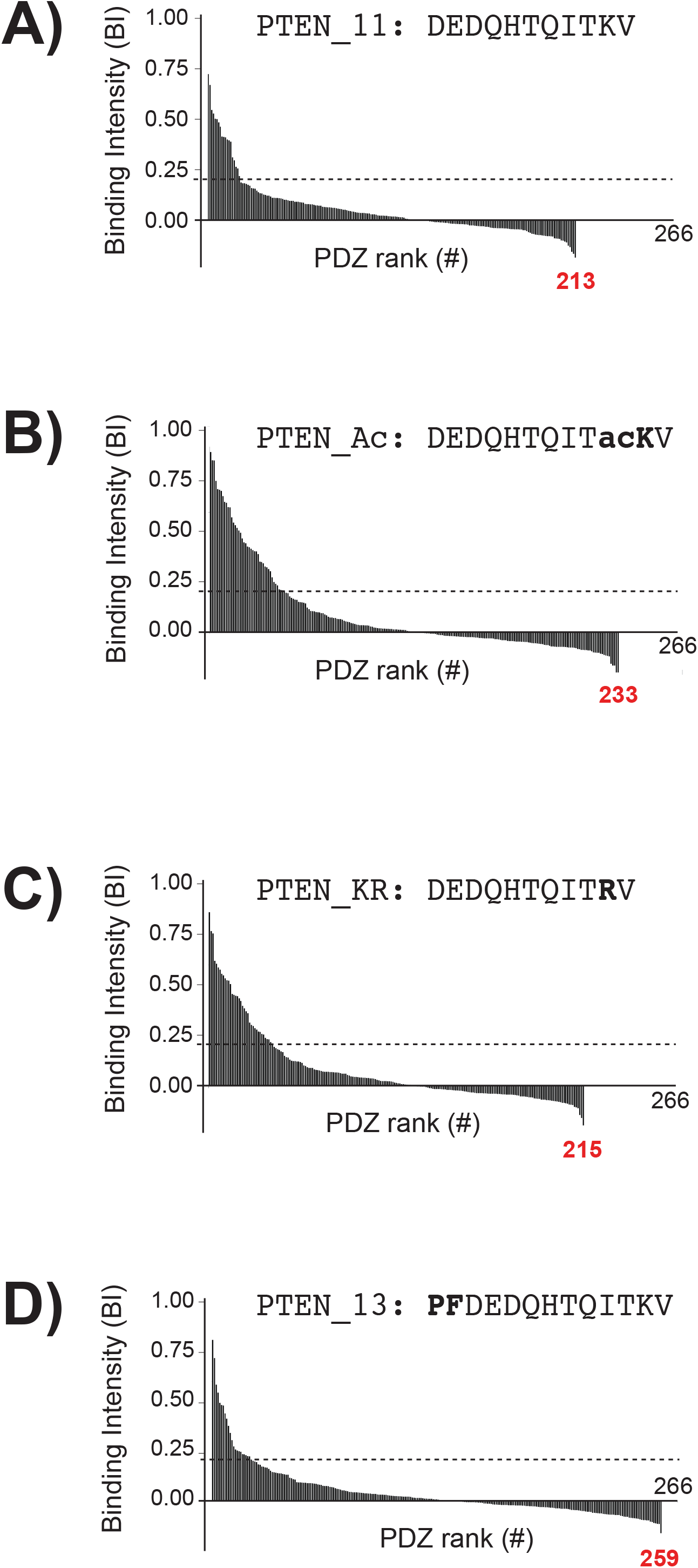
PDZ binding profiles of the four PTEN peptides. **Holdup** binding profiles obtained are shown for PTEN_11 (**A**), PTEN_Ac (**B**), PTEN_KR (**C**) and PTEN_13 (**D**). In each profile, the PDZ binders are ranked from left to right of the plot in BI decreasing order along the X-axis. Data for all the measured holdup data are shown. The grey dotted line shows the threshold for confidence value, set at BI = 0.20 (see main text). For each experiment, the number of PDZ domains for which we obtained a measurement that passed the quality filtering step, and could therefore be included in the plot, is indicated (red case numbers). The holdup data for PDZ / PBM pairs with BI>0.20 are shown in Supp. Fig. S1. *Single column fitting image.*

### Orthogonal validation by competitive FP and conversion of holdup BI data into dissociation constants of the four PTEN PBMs versus the human PDZome

Calculation of an equilibrium constant for a PDZ-PBM interaction requires three concentrations: free PBM, free PDZ and PDZ-PBM complex. The holdup assay delivers for each PDZ / PBM pair the concentrations of free PDZ and PDZ-PBM complex, but not that of free PBM. To circumvent this problem we systematically measured by competitive FP, an orthogonal approach to holdup, the K_D_ constants for the 4 PTEN peptides against a subset of 20 PDZ domains (**Supp. Fig. S2**), resulting in approx. 8 to 10 significant K_D_ for each PBM. These accurate dissociation constants were used to back-calculate the peptide concentrations in the holdup assays (**Fig. 4A**). We found the concentrations of the different PBM peptides to vary between 10 and 90 μM, with averages between 17 and 34 μM depending on the PBM after outlier rejection. A global mean of 26 μM considering all the peptides was determined. A plot of experimental K_D_ obtained by FP *versus* BI superimposed well with the theoretical affinity values calculated using the global average peptide concentration of 26 μM (**Supp. Fig. S3**). This shows a very good agreement between the holdup BI values and the binding strength derived from competitive FP measurements.

**Fig 4.**
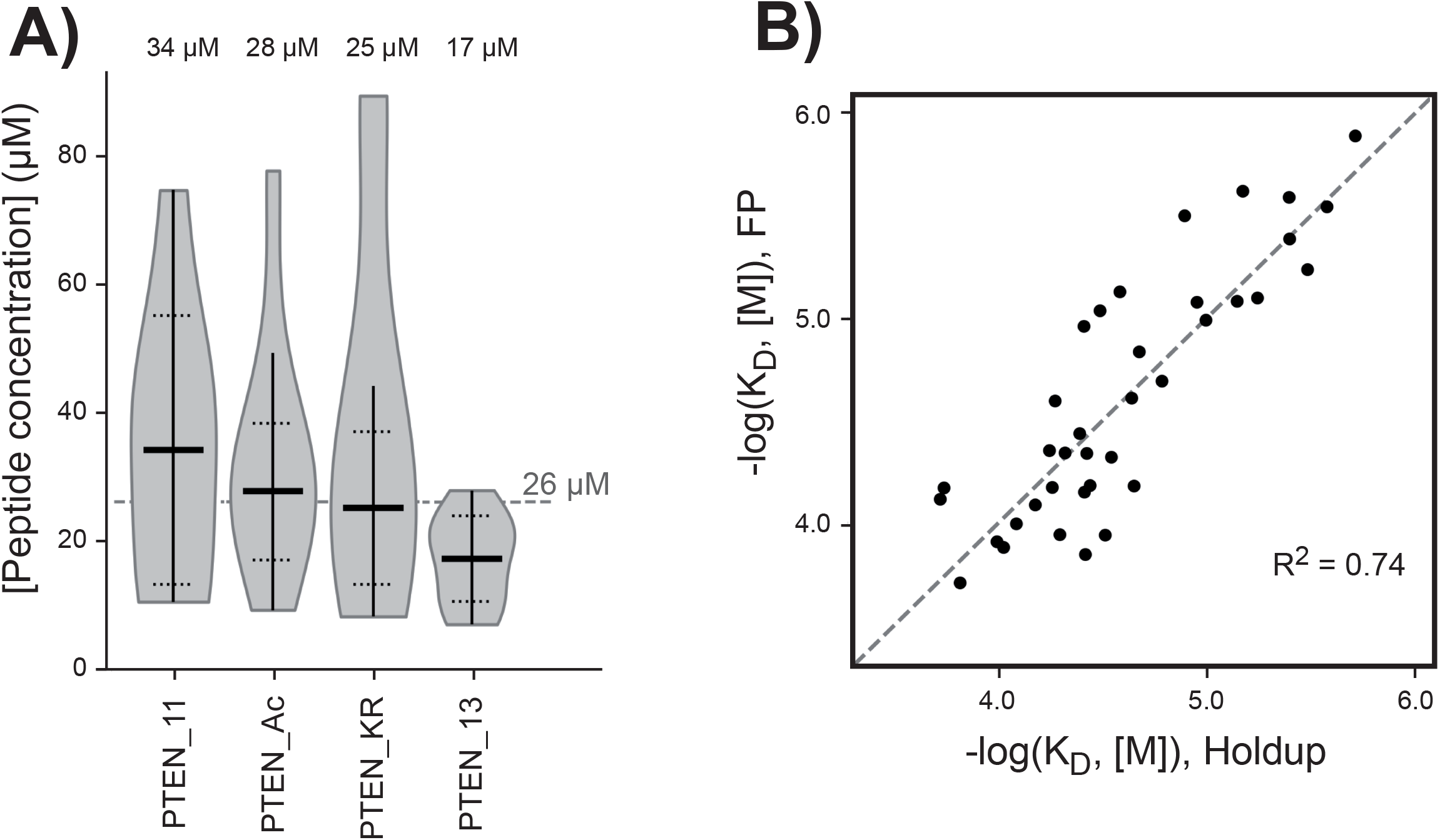
Conversion of the holdup binding intensities into affinities constants. (**A**) The violin plots shows the distribution of all the back-calculated apparent peptide concentrations obtained when both a quantifiable and significant (>0.20) BI value by holdup and a dissociation constant by FP were available for a given PDZ / PBM pair. The vertical line indicates the range of the distribution while the horizontal lines show the final mean peptide concentration and its final standard deviation after outlier exclusion (considering the 3σ rule). The final average peptide concentrations represented by the thick lines are used to convert the holdup BI values into K_D_. (**B**) Comparison between the converted dissociation constants from the holdup assay and the dissociation constants directly measured by FP assay. The dotted line represents the perfect theoretical correlation. Since the data points seem to be randomly distributed on both sides of this dotted line, the R^2^ is indicative of the goodness of fit. *Single column fitting image.*

Using the mean concentration obtained above for every PTEN peptide, the experimental BI values recorded by holdup for all tested domain / peptide pairs were subsequently transformed into equilibrium dissociation constants. A strong agreement is observed between the affinity constants obtained from holdup and FP assays with a coefficient of determination R^2^ = 0.74 (**Fig. 4B**), confirming that singlicate holdup runs provided highly reliable data. At this stage, affinity data measured by FP assay were also included for the few PDZ domains (MAST1-1, MAST2-1, SNX27-1, MAGI1-2 and GRID2IP-2) for which holdup data were missing according to the quality score filtering, representing 1 to 3 additional PDZ binders per PTEN construct. A total of 215, 234, 218 and 259 interaction data were obtained for PTEN_11, PTEN_Ac, PTEN_KR and PTEN_13, respectively. The transformation into affinity values makes then possible to compare binding affinity profiles obtained for different peptides and different batches.

### From binding profiles to specificity quantification

The above described holdup-FP strategy delivers binding affinity constants, a universal chemical property. The affinity values obtained for each PTEN peptide were plotted in a logarithmic scale, hence proportional to free energies of binding ΔG at a fixed temperature (**Fig. 5**). The resulting profiles contains information about specificity or promiscuity since a promiscuous peptide as seen by hodlup would bind to a large number of PDZ. We then looked for a numerical parameter that would express, in a quantitative way, this specificity or promiscuity information. For this purpose, we calculated for each profile the difference between the maximal and minimal affinity values detected by the assay, ΔG_max_ − ΔG_min_. Next, we introduced a threshold affinity, called “half-maximal binding affinity” defined as follows: ΔG_half_ = ΔG_min_ + (ΔG_max_ − ΔG_min_)/2. We then defined the half-maximal binding promiscuity index I_P_ as the percentage of PDZ domains bound to the PBM with an affinity superior to the half-maximal affinity relative to the total number of PDZ domains that were successfully measured in the assay (**Fig 5**). Alternatively, the specificity index I_S_ could be defined as 1 – I_P_. Therefore, the lower the promiscuity index, the higher the specificity index, the higher the specificity of the PBM for a few selected domains across the PDZome. For instance, if 250 PDZ domains were fully assayed, and only 5 PDZ domains bound to the PBM with an affinity superior to the half-maximal affinity, the specificity index will be 98%. If 25 domains bound with an affinity superior to the half-maximal affinity, the specificity index will be 90%.

**Fig 5.**
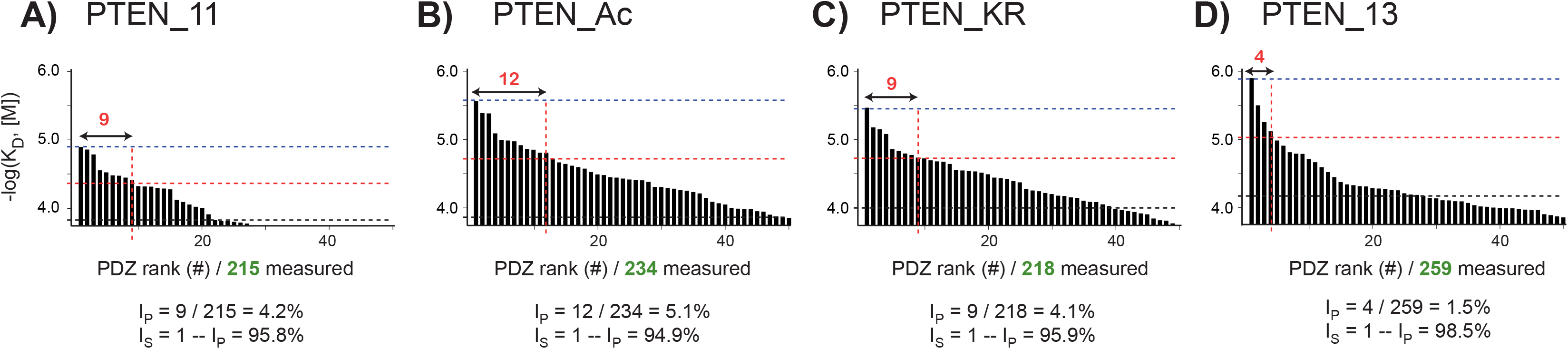
Determination of the specificity index for the PTEN binding profiles. For every profile, the significant PDZ binder affinity values are ranked from left to right along the X-axis in -log(*K*_D_) decreasing order. The non-significant or undetected binders were omitted for clarity. The grey dotted line corresponds to the threshold BI value after converting it into −log(*K*_*D*_) scale, while the blue and red dotted lines represent the highest affinity and the affinity at half the difference between the maximal and weakest significant affinity values, respectively. The reader can note that, for a constant threshold BI value (0.20), the weakest affinity values may vary moderately due to non-constant peptide concentrations. The numbers of PDZ domains above the half-maximal binding affinity” are indicated in red, while the numbers of tested and validated PDZ domains are in green. Values calculated for the promiscuity index (I_P_) and the specificity index (I_S_) are given (see main text). Full data sets for holdup and FP are visible in **Supp. Fig. S1 and S2,** respectively. *Double column fitting image.*

We probed the specificity index on the PDZome-binding profiles of the four PTEN peptides. In both the BI-based and the affinity-based representations (**Fig. 2 and 5**), the shapes of the profiles of PTEN_11, PTEN_Ac, PTEN_KR look similar, while the PTEN_13 presents a sharper, faster-decreasing profile. This indicates, in qualitative terms, that the PTEN_13 peptide selects PDZ domains in a more specific-less promiscuous-way than that of the three of other peptides. This is fully confirmed by the computed specificity indexes, which yield close values for PTEN_11, PTEN_Ac, and PTEN_KR (95.8%, 94.9%, 95.9%, respectively), while the extended wild-type peptide PTEN_13 displays a higher specificity index (98.5%) indicative of a higher specificity towards a few selected PDZ domains.

### Rearrangements of the binding profiles due to minor changes in PTEN

The PTEN-bound PDZ domains are distributed over a diversity of PDZ-containing proteins (**Fig. 6**). Several PDZ domains such as MAST2-1, PDZD7-3, SNX27-1, MAGI1-3 and GRASP-1 were systematically among the strongest interaction partners of all four PTEN PBM variants. We compared our data to previously published studies, bearing in mind that sequences and boundaries of PTEN and PDZ constructs may differ (**Table 1**). Our results agree with isothermal titration calorimetry data obtained for SNX27-1 / PTEN [38] and MAST2-1 / PTEN complexes [26] and, in part, with FP data obtained for PARD3-1 / PTEN complex [39]. Interestingly, some of our newly identified PTEN-binding PDZ domains, such as MAGI1-3, MAGI2-3 and DLG4-1 bound wild-type PTEN peptides with a stronger affinity than the domains of the same proteins that were previously published to bind PTEN, such as MAGI1-2 [40], MAGI2-2 [24] and DLG4-3 [41], respectively. This result illustrates the strength of the complementary holdup / FP approach which can provide an affinity ranking of PDZ domains even within multi PDZ-containing proteins.

**Fig 6.**
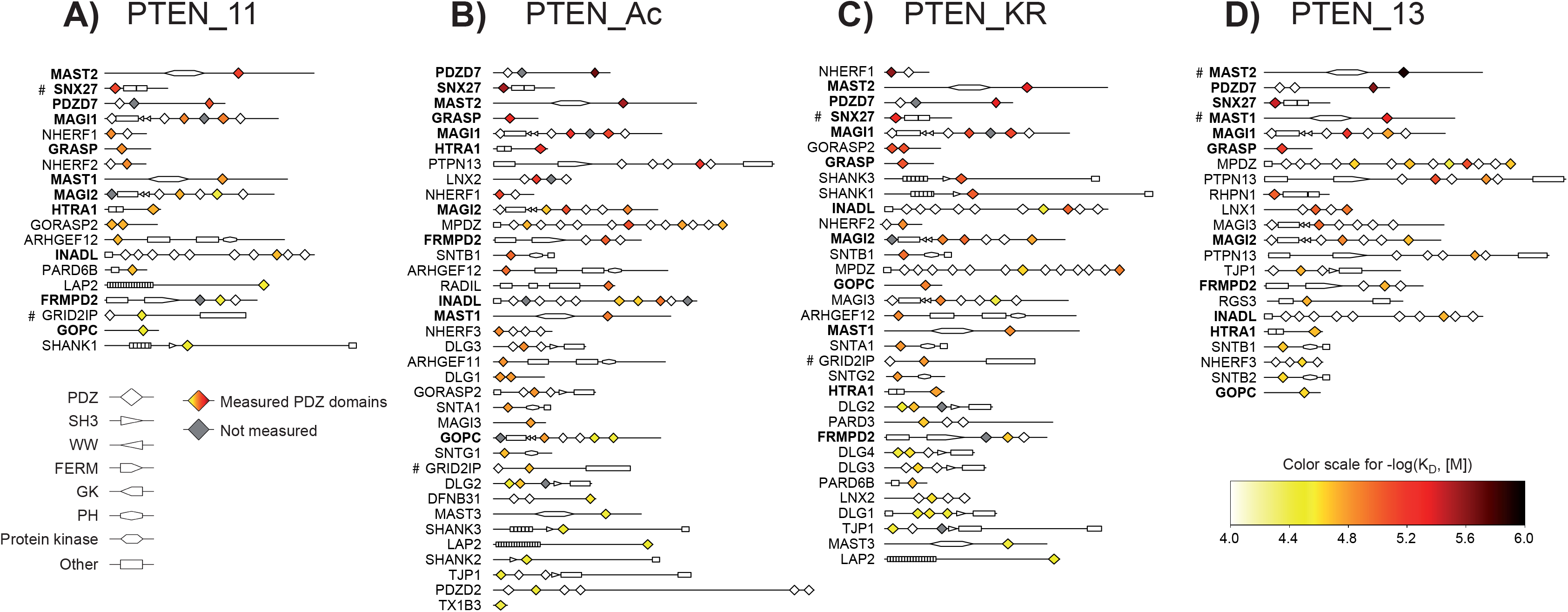
Domain representations of the impacted PDZ domains by the different PTEN peptides. Proteins containing PDZ domains significantly bound to one PTEN peptide are colored and ranked from strongest to weakest binding strength depending on the best individual PDZ binder within each protein. The color code from white to black is indicative of the −log(*K*_*D*_) values in the range of 4.0 – 6.0 after filtering step and BI conversion. The symbol (#) denotes PDZ domain for which the BI value could not be measured directly by holdup and has been inferred from FP measurements. Protein names appeared in bold when significant −log(K_D_) values are observed for the four PTEN PBM. *Double column fitting image.*

**Table 1.**
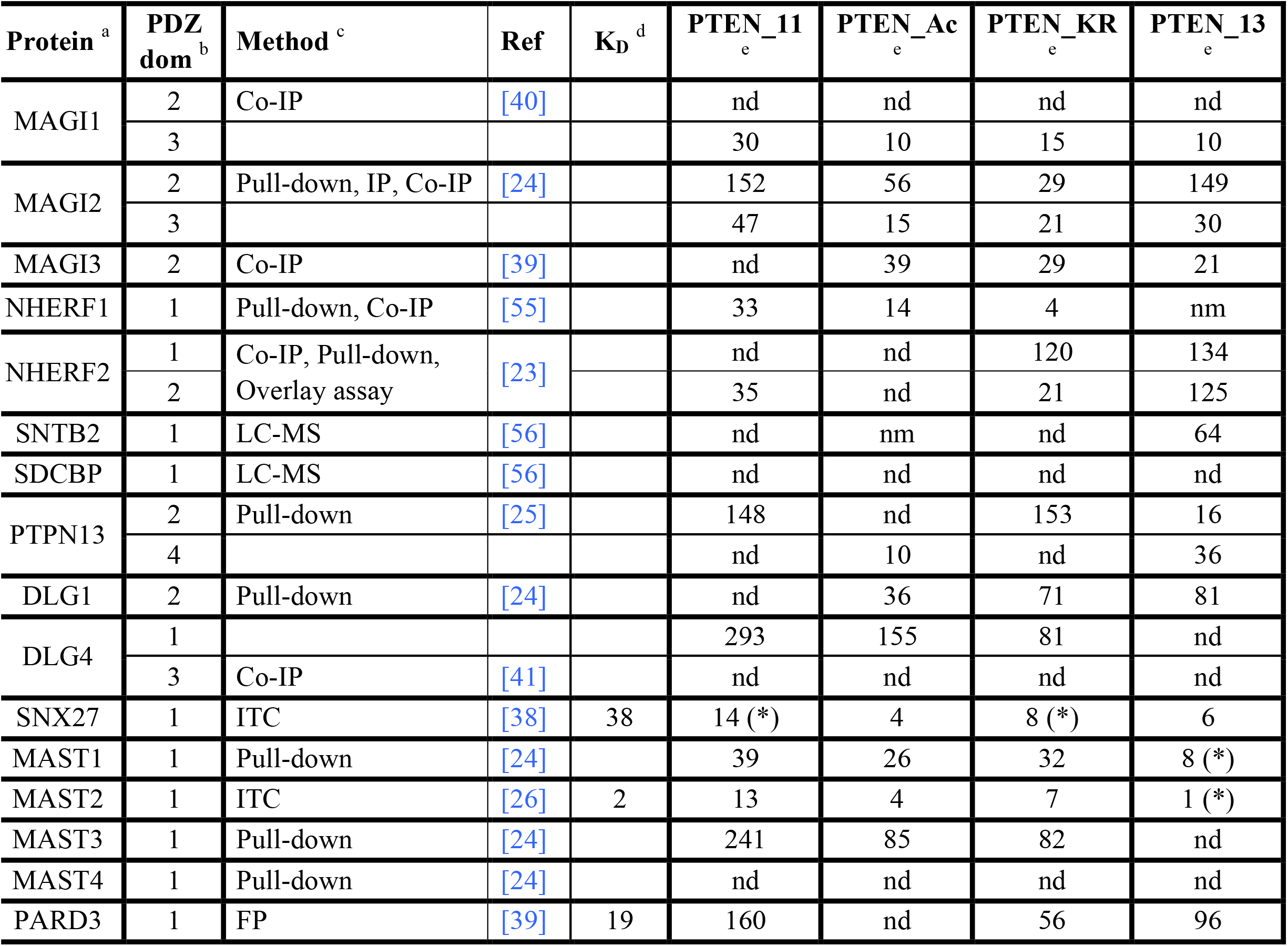
PDZ domains interactors for PTEN according to literature and the present study. Each row corresponds to a protein for which a binding to PTEN has been been described in literature. The main methods and the PDZ domain number are indicated. The four last columns contain information obtained by combining the holdup and FP methods in the present study. ^a^ Protein name ^b^ Domain interaction site for PTEN ^c^ Detection methods described in literature ^d^ Affinity provided in the literature when available (in μM) ^e^ Affinity measured by holdup in this study (in μM) * Affinity measured by FP in this study (in μM) IP: Immunoprecipitation Co-IP: Co-immunoprecipitation nd: not detected in the holdup assay nm: not measured in the holdup assay

Although the shapes of the dissociation constant profiles for the three 11-mer PTEN variants were globally similar, the PDZ domains are reshuffled between the various profiles (**Fig 7**). We detected at least 20 additional new partners for PTEN_Ac, and 11 for PTEN_KR (**Fig. 7A & Supp. Info. S1**). The acetylated peptide is highly promiscuous and binds to all the partners of the native PTEN_11 PBM, plus numerous additional ones. Furthermore, the arginine mutation does not seem able to efficiently reproduce the acetylated state as seen by the number of partners (8 over a total of 37) detected for PTEN_KR and not for PTEN_Ac. The opposite effect with 15 over a total of 43 detected for PTEN_Ac and not for PTEN_KR is even more pronounced, suggesting that the acetylation effect on binding is mainly due to the acetyl group rather than the size of the side chain carried by the acetylated lysine residue.

**Fig 7.**
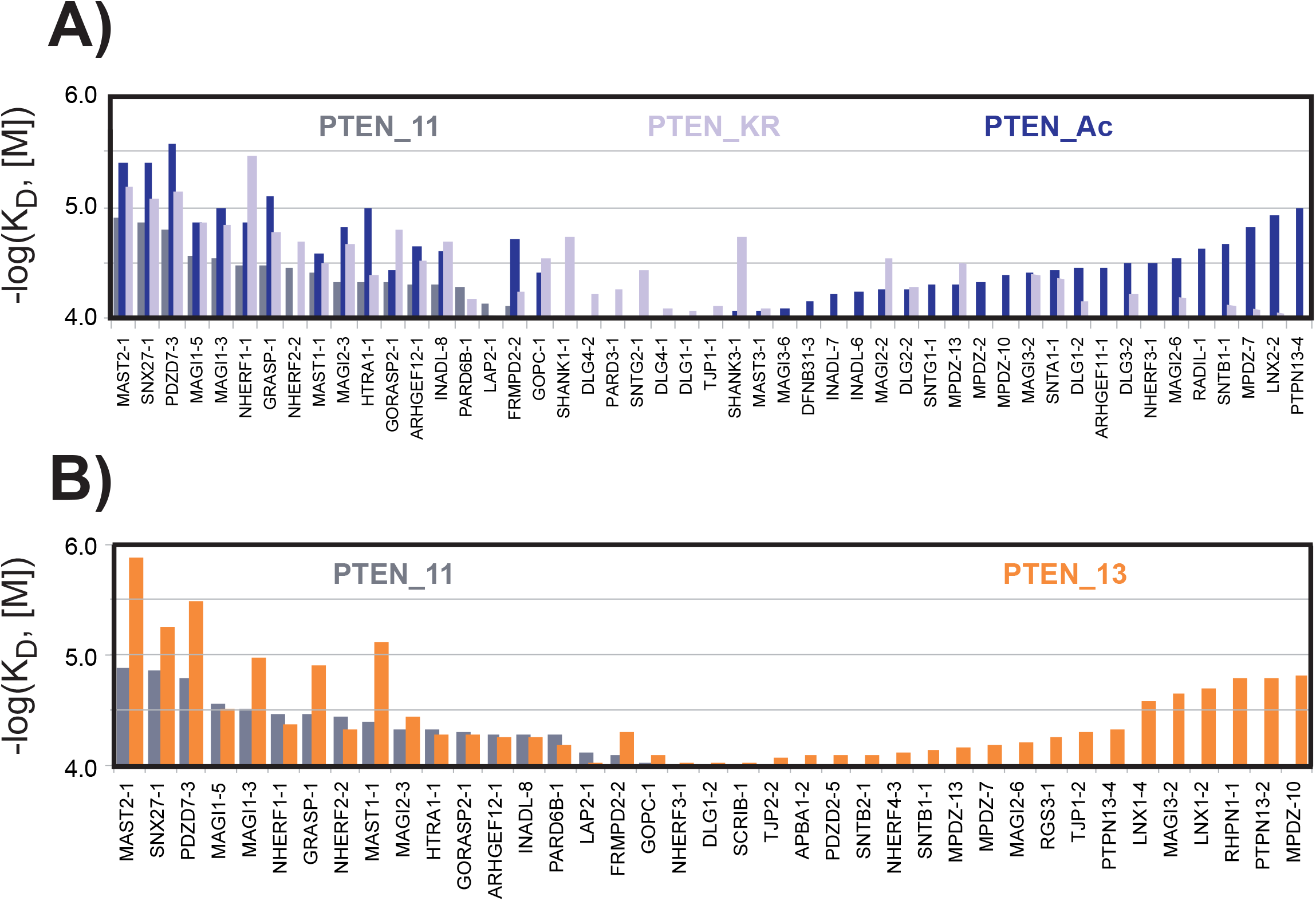
Changes in the PDZ binding profiles induced by changes in the PTEN peptides. (**A**) Comparison between PTEN_11 (grey), PTEN_KR (light purple) and PTEN_Ac (dark blue) using a shared PDZ axis. For the wild-type PTEN_11 peptide, the PDZ domains were ranked in descending affinity order along the X-axis, from left to right according to the significant affinities for PTEN_11, and from right to left according to the significant affinities solely detected for PTEN_13. The remaining PDZ domains that bind only to the PTEN_KR peptide were added in the middle region. (**B**) Comparison between PTEN_11 (grey) and PTEN_13 (orange) on a shared PDZ axis. The PDZ domains were ranked along the X-axis in descending order, from left to right according to the significant affinities for PTEN_11, and from right to left according to the significant affinities exclusively detected for PTEN_13. The left and right regions thus show PDZ domains that prefer the shorter or the longer PTEN PBM version, respectively. The overall uncertainty on log(*K*_*D*_) values was estimated to be roughly ± 0.2 in log(M) unit by propagating BI uncertainty estimated in previous studies. *Double column fitting image.*

The impact of the PTEN peptide length was noticeable by comparing the dissociation constant profiles of PTEN_11 and PTEN_13 (**Fig. 7B**). The detected interactions of PTEN_13 were markedly stronger compared to the affinities observed for the same PDZ partners in PTEN_11. The strongest effect is observed for MAST2, the top binder for both PTEN_11 and PTEN_13, for which the −log(K_D_) value increases from 4.9 to 5.9 in log(M) unit (i.e. a jump from 13 μM to 1 μM), corresponding to about a 10 fold stronger affinity. Only a few interactions, in the low range affinities, were potentially slightly strengthened although most likely not significantly. Moreover, 24 new binders appear due apparently to the presence of the two extra residues in the N-terminus of the peptide. These rearrangements are particularly noteworthy since the mutations or the Pro-Phe inclusion introduced for this work are located at positions described as non-critical for PBM binding.

## Discussion

### Insight into the holdup: a powerful semi-automated tool for medium-to-low affinity measurements

In this work, we quantitatively assessed more than 1,000 distinct PDZ-peptide affinities by using a “crude holdup assay” protocol, which quantifies the disappearance of a single protein peak (the tag-PDZ peak) out of a complex crude overexpression extract. This protocol requires a rigorous approach. Some critical biochemical steps have been previously identified [28] [29] including the standardized expression of the complete PDZome, the verification of its quality, the calibration of its concentrations in the crude extract, and a careful quality control of capillary electrophoresis runs. For data treatment, we developed a computational processing step for accurate superimposition of the electropherograms to improve the precision of binding intensities [33]. Here, a four-criteria quality score was introduced to further rationalize data curation. These improvements allow us to minimize the amount of false positive and false negative results. In addition, to spare costs and manpower for data treatment, holdup experiments were run in singlicate and combined with an orthogonal approach, the competitive FP. This generated high-confidence affinity data and allowed us to convert holdup binding intensities (BI) values into affinities (ΔG or *K*_*D*_). The use of such an intrinsic universal parameter of molecular complexes also presented the advantage to facilitate the comparison with data available in the literature. In future developments of the automated holdup assay, we envision to replace crude overexpression extracts by purified proteins, which greatly facilitate both readout and data treatment [31].

### Impact of PTEN PBM acetylation on its PDZ interactome

Lysine acetylation is a PTM difficult to study and reproduce *in vitro*. Some studies have explored lysine acetylation by proteomic approaches [42], while others have mutated lysine residues to glutamine or arginine to mimic acetylation or suppress acetylatability, respectively [21],[43]–[45]. In the present study, we investigated with chemically synthetized peptides that allow to fully control PTM the differential effects of acetylation or mutation of a lysine residue on the PDZ interactome of PTEN. PTEN is a tumor suppressor that is frequently inactivated in human cancers [46],[47]. Some *in vivo* activities of PTEN such as PI3K signaling regulation seem to be abolished when PTEN is acetylated [48]. In addition, the Lys-to-Arg mutation at PTEN position 402 (corresponding to non-essential p-1 position of its C-terminal PBM) abolished PTEN acetylatability [21]. However, this may either mean that K402 is a direct acetylation target or indicates that the integrity of the PTEN PBM sequence is required for PBM-dependent acetylation of PTEN at other sites distinct from K402.

We found that K402 acetylation (inducing a loss of a positive charge and a slight increase of bulkiness) altered both the strength and the number of detected PDZ binders of PTEN. In contrast, the K402R mutation (preserving the positive charge but further increasing the bulkiness) did not alter the overall binding strength nor the number of binders. Furthermore, the K402R mutant retains binding to most partners of the native motif and also binds to a subset of the acetylated peptide partners. Therefore, at the p-1 position of the PTEN PBM, the presence or absence of a positive charge appears more critical for PDZ recognition than the bulkiness of the side chain.

Although a few PDZ domains including several from MAGI and NHERF detectably bound to all three peptides PTEN_11, PTEN_Ac and PTEN_KR, several PDZ domains bound only one or two of those peptides. For instance, both PTEN_Ac and PTEN_KR bound stronger than wild-type PTEN_11 to MAGI2_2 or DLG1_2 domains, in agreement with Ikenoue *et al*. Since our study is performed over the full PDZome, this implies that acetylation generally increases the affinity of PTEN for PDZ domains. Overall, the rather large number of PDZ partners associating with the PTEN PBM confirms that domain / motif networks are rather promiscuous [49].

### Lessons from distal residues on the PTEN interactome

There is no consensus for the precise residue length of a given PBM needed to complete the interaction with a PDZ domain. Although the four C-terminal residues are usually thought to constitute the core of a PBM, it was shown that peptides comprising the last 10 positions of a PBM undergo a significant change in their PDZ-binding affinities as compared to peptides comprising only the last 5 positions [13]. Such affinity variations may result from differences of entropy of the free peptides, from altered interface contacts in the resulting PDZ-PBM complexes, or a combination of both. Accordingly, synthetic or recombinant PBMs employed for PDZ interactions generally include at least 9 to 11 residues [4],[5],[15],[17],[28],[50]. Indeed, the presence of distal sites altering PDZ-PBM binding has already been described [51], even at positions as far as at p-36 [52]. In the particular case of PTEN, Terrien *et al*. previously demonstrated the existence of a distal “exosite” at F392 (p-11), that triggers novel contacts within a secondary exposed hydrophobic surface of MAST2 [26]. Here, we showed that the inclusion of two extra residues, including F392, (PTEN_13 versus PTEN_11) affected both the PDZ interactome identified for PTEN and the specificity of its PBM. Indeed, several PDZ domains detectably bound only to the longer construct, in line with the idea of a global affinity increase because of the larger number of atomic contacts. Furthermore, while the three 11-mer peptides displayed equivalent PDZ-binding specificity, PTEN_13 showed an increased specificity. The addition of the distal exosite was therefore more influential for specificity than the chemical variations (Lys acetylation or Lys to Arg mutation) at p-1 position.

In principle, one may argue that domain-motif binding events may be altered by any distal region, so that only studies full-length protein / protein interactions are relevant. Notwithstanding the methodological issues (large full-length proteins can be very difficult to handle), one must keep in mind that most full-length multi-domain proteins are prone to many conformational changes (inducible by partner binding, ligand binding, PTM, molecular crowding, and so forth), which in turn influence the availability of their globular domains or linear motifs for binding events. This justifies the ‘domainomics’ approach [53] undertaken in this work, that focuses on the binding properties of minimal interacting fragments of proteins, such as a globular domains (e.g., PDZs) and short linear motifs (e.g., PBMs). Even if our binder list might be incomplete as compared to studies involving full-length proteins, it provides a list of the PDZ domains capable to interact with the motif of the PTEN PBM, constituting the minimal block at the binding interface of protein / protein interaction.

### To bind or not to bind

In this work, by covering almost the entire PDZ family, we quantified both the number of interacting and non-interacting partners for a given PBM. The knowledge of the two numbers is important since the count of 3 binding partners over a dataset of 10 domains, or 3 partners over a dataset of 100, is not reporting the same specificity. Over the years, we have accumulated holdup data for many peptides and noticed that more than 90% (244/266) of the PDZs in our expressed PDZome are functionally active since they interacted significantly with at least one PBM [29]. This indicates that most of the non-binders detected in our profiles are trustable. The holdup assay is therefore a reliable approach to address not only the specificity but also the ‘negatome’ in the sense of the negative interaction dataset as originally proposed [54].

In this work, we derived from the PDZome-binding profiles a single numerical index to evaluate the degree of specificity of a given PBM towards particular PDZ domains. One can assume the probability of binder occurrence to be all the more similar in the validated and untested PDZ datasets as the validated dataset is covering a large part (>~80%) of the entire human PDZome. The calculation of the specificity index will thus be roughly the same for both the validated and the complete PDZ datasets. One must notice that this index is not fully satisfying and cannot be considered as a universal parameter beyond our particular PBM-PDZome affinity profiling studies. In particular this index is only operative to compare profiles with a roughly continuous decreasing shape, e.g. in absence of discontinuous “breaks” or “stairs”. But the concept of specificity index affords the advantage of introducing a numerical value attached to each PBM profile, that will ease their comparison.

## Conclusion

In this study, we showed that the hydrophobic exosite at position p-11, not only impacts the interaction of the PTEN C-terminal tail with MAST2 as previously reported [26], but also affects its binding to a large set of other PDZ interaction partners, suggesting to well control the length of the polypeptide used for *in vitro* interaction studies. More importantly, we also showed that both, the K402 acetylation and even the K402R point mutation at p-1, a non-critical position of the canonical PBM motif for PDZ / PBM interaction, significantly increased the number of targeted PDZ domains. This could be of primary relevance, knowing that the activities of the tumor suppressor PTEN protein is regulated by acetylation. Finally, we also introduced a way to quantify specificity that could be extended to other interaction studies covering a whole domain family.

## Abbreviations

PDZ: PSD95/DLG/ZO-1
PBM: PDZ-binding motif
HPV: Human Papilloma Virus
FP: fluorescence polarization
MBP: Maltose-Binding Protein
TRX: Thioredoxin

## Supplementary information

– **Supp. Fig. S1**: contains the entire data set obtained by holdup for BI>0.20. For each panel, after superimposition of the two electropherograms recorded for the PBM of interest (blue dotted line) and for the biotin reference (black solid line), the normalization of the electropherogram of the PBM compared to the one of the reference is done using the signal of the lysozyme added in every sample at a constant concentration (red peak). The region between 20 and 60 kDa which contains peaks of the crude extract supposedly to be constant, is used to verify the proper intensity normalization of the two electropherograms. The intensities of the peak of interest after proper alignment along the molecular weight scale (region covered by the green dotted line) are subsequently used to quantify the depletion of an individual PDZ domain and then the BI value. All those normalization and alignment steps are performed automatically.
– **Supp. Fig. S2**: contains the entire data set obtained by FP. Average of FP data recorded in triplicate are represented with black dots. The reported dissociation constants and errors are the average and the standard deviations of 500 independent Monte-Carlo simulations, calculated using ProFit as described in Simon et al., 2020.
– **Supp. Fig. S3**: contains the experimental (BI, K_D_) plot superimposed with K_D_ obtained with **Eq. 2**. Error bars are representative of peptide concentration uncertainty after propagation to the −log(K_D_) values.
– **Supp. Info S1** file: contains the data set with all the BI values together with the transformed dissociation equilibrium constants for each PDZ-PBM interaction. All the plots in this study are performed according to this data set.

## Acknowledgements

This work received institutional support from Centre National de la Recherche Scientifique (CNRS), Université de Strasbourg, Institut National de la Santé et de la Recherche Médicale (INSERM) and Région Alsace. G.G. was supported by the “Post-doctorants en France” program of the ARC foundation. The work was supported by funding from the European Union’s Horizon 2020 research and innovation program under the Marie Sklodowska-Curie grant agreement No 675341, by the Ligue contre le cancer (équipe labellisée 2015), by the National Institutes of Health (Grant R01CA134737), by the Canceropôle Grand-Est (projet Emergent), and by the French Infrastructure for Integrated Structural Biology (FRISBI).

## Author contributions

PJ: conceptualization; data analysis and interpretation; drafting and revising the article. GG: conception and design; data acquisition; data analysis and interpretation; drafting and revising the article. CK: data acquisition; revising the article. GB: data analysis; revising the article. VG: data acquisition; revising the article. CCS: data acquisition; revising the article. PE: peptide synthesis; revising the article. RV: data acquisition, revising the article. NW: funding acquisition; conception and design; data analysis and interpretation; revising the article. GT: funding acquisition; conception and design; data analysis and interpretation; drafting and revising the article. YN: funding acquisition; conception and design; data analysis and interpretation; drafting and revising the article.

