## Supplementary figures and images for "Interactomic affinity profiling by holdup assay: acetylation and distal residues impact the PDZome-binding specificity of PTEN phosphatase"

### Supp. Fig. S1

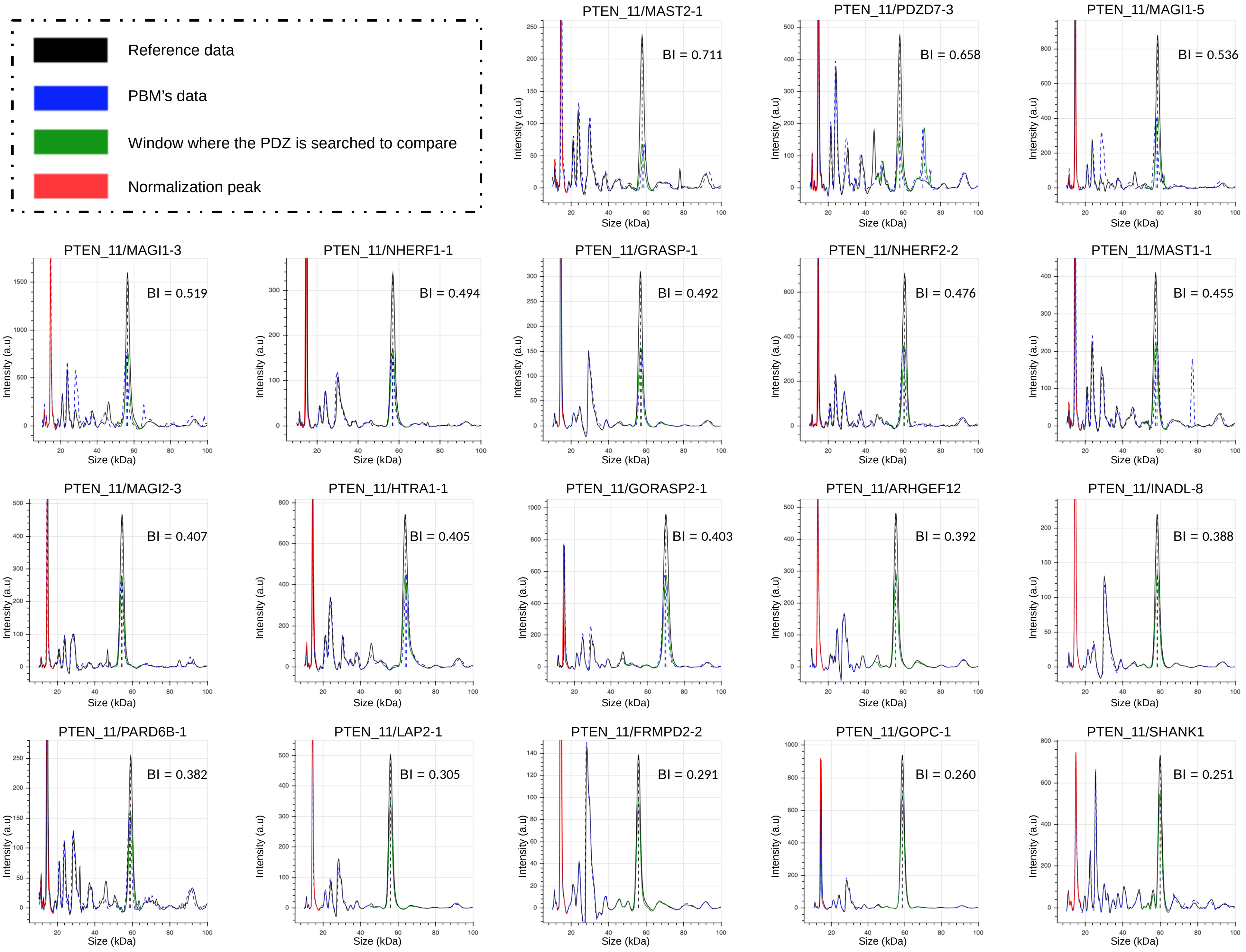

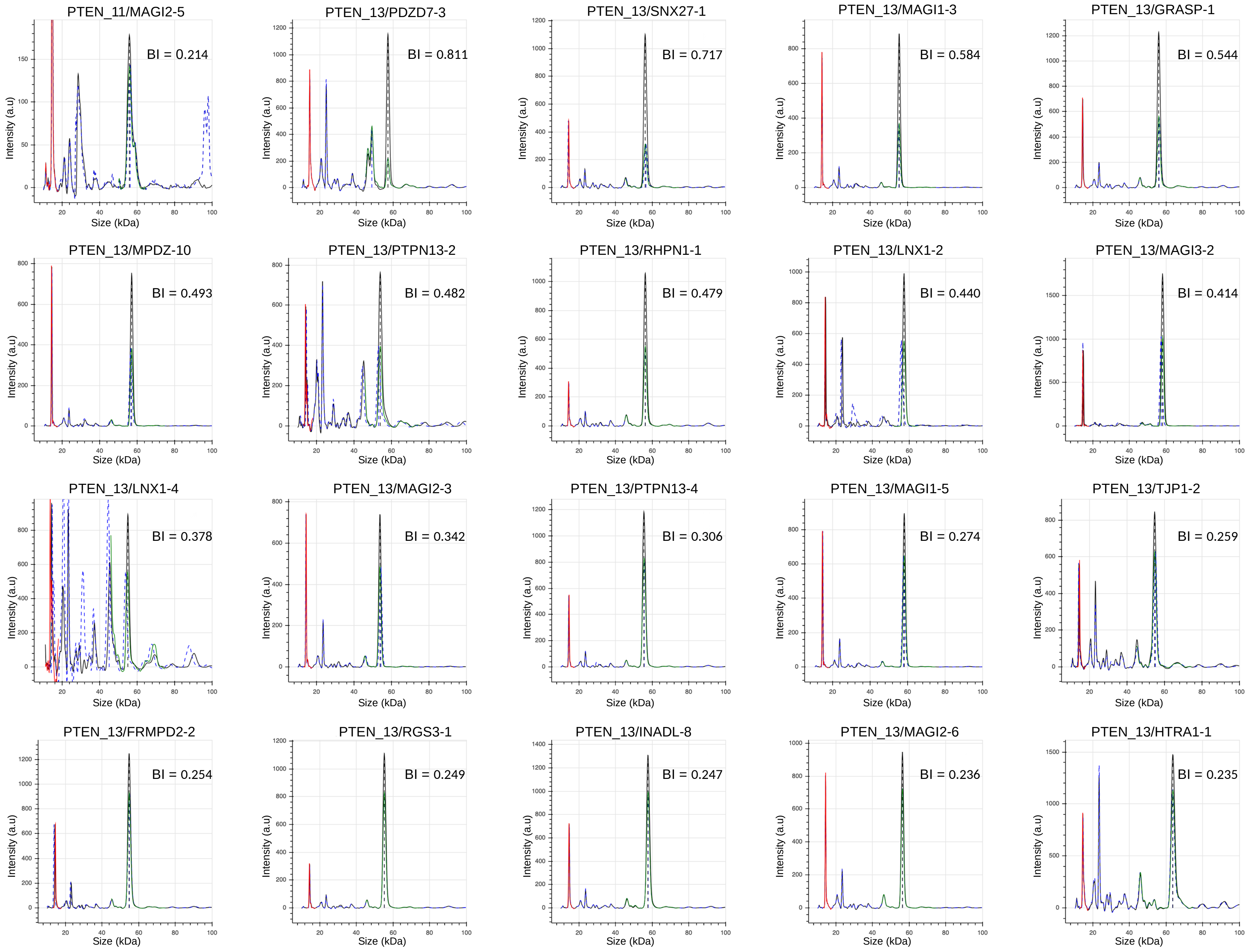

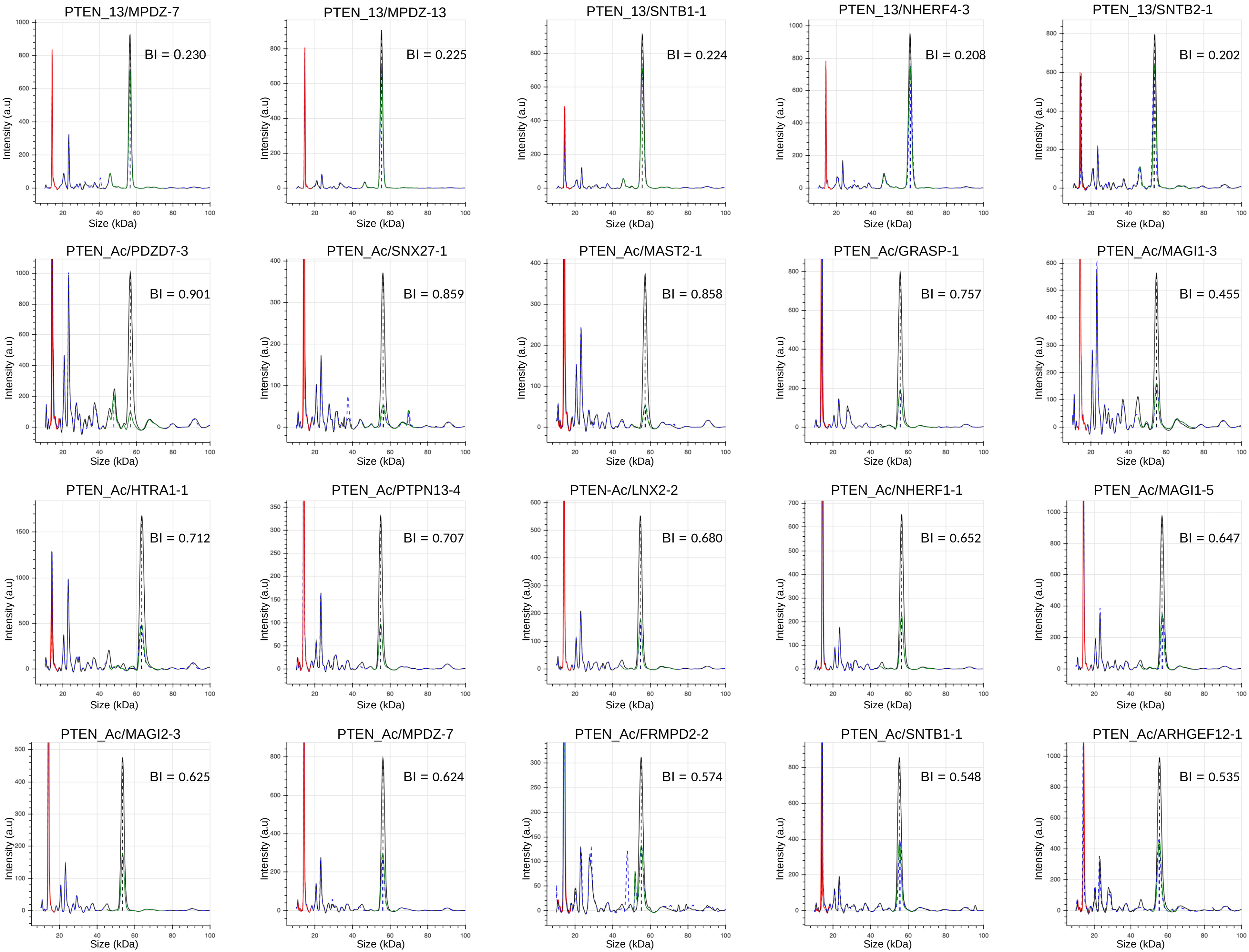

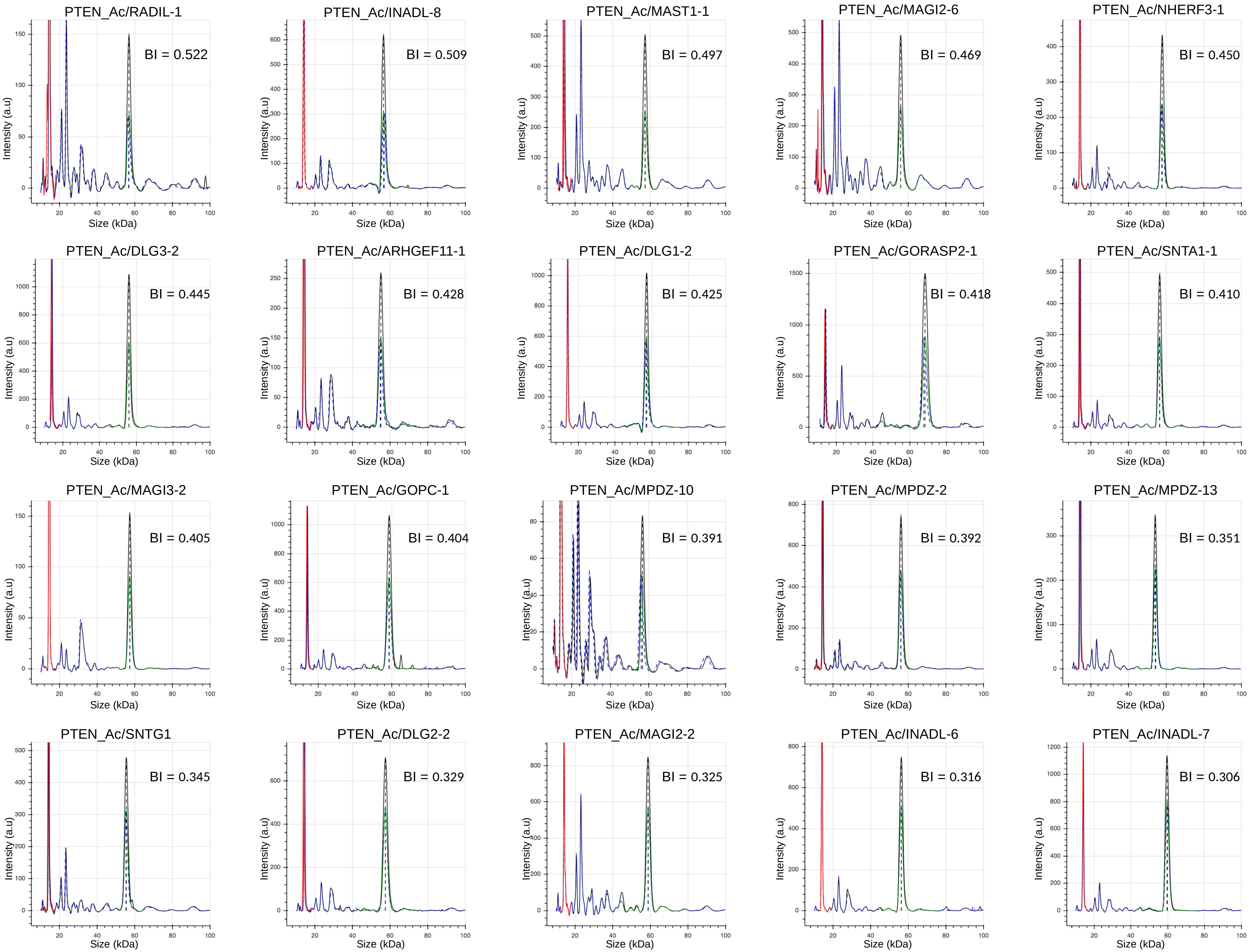

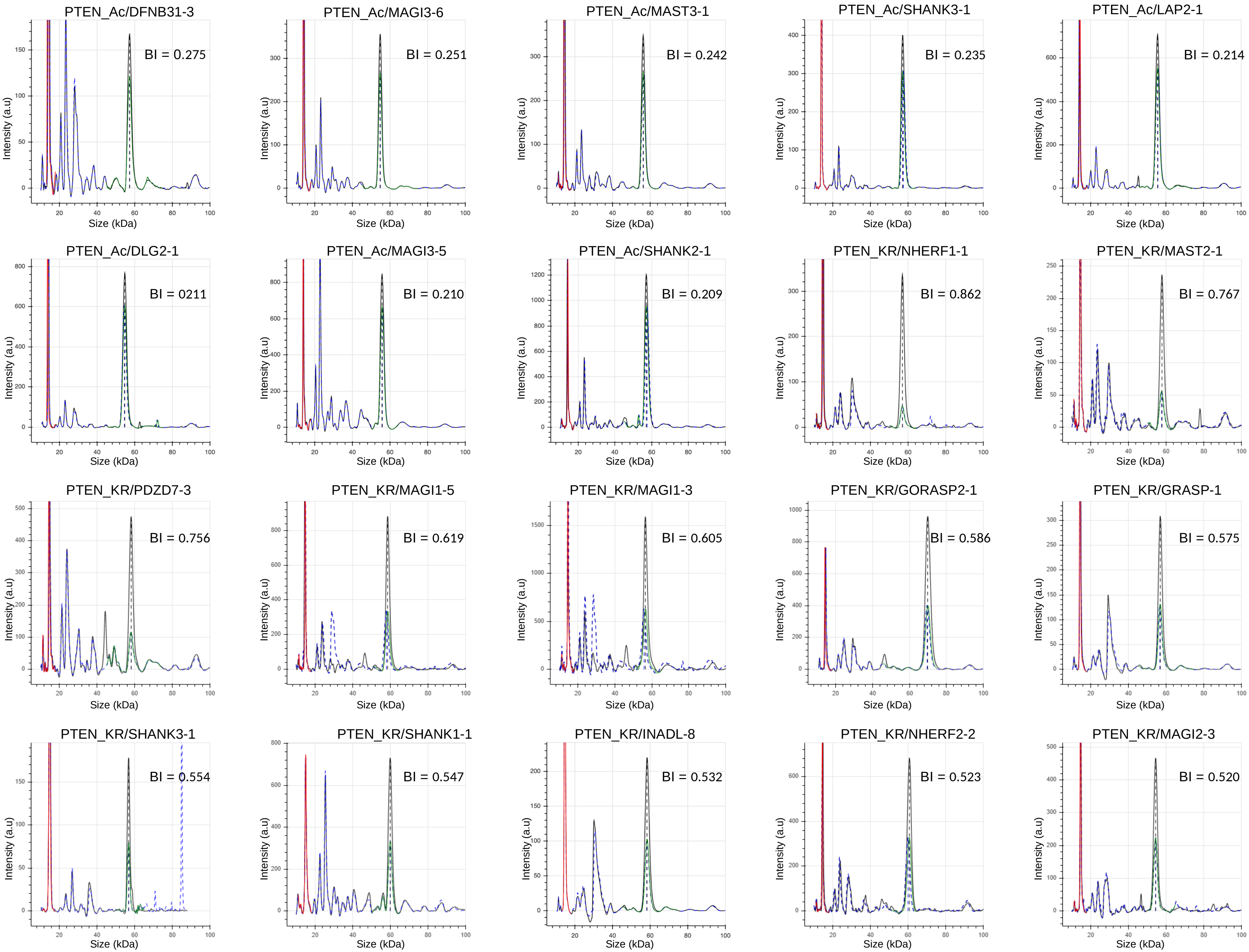

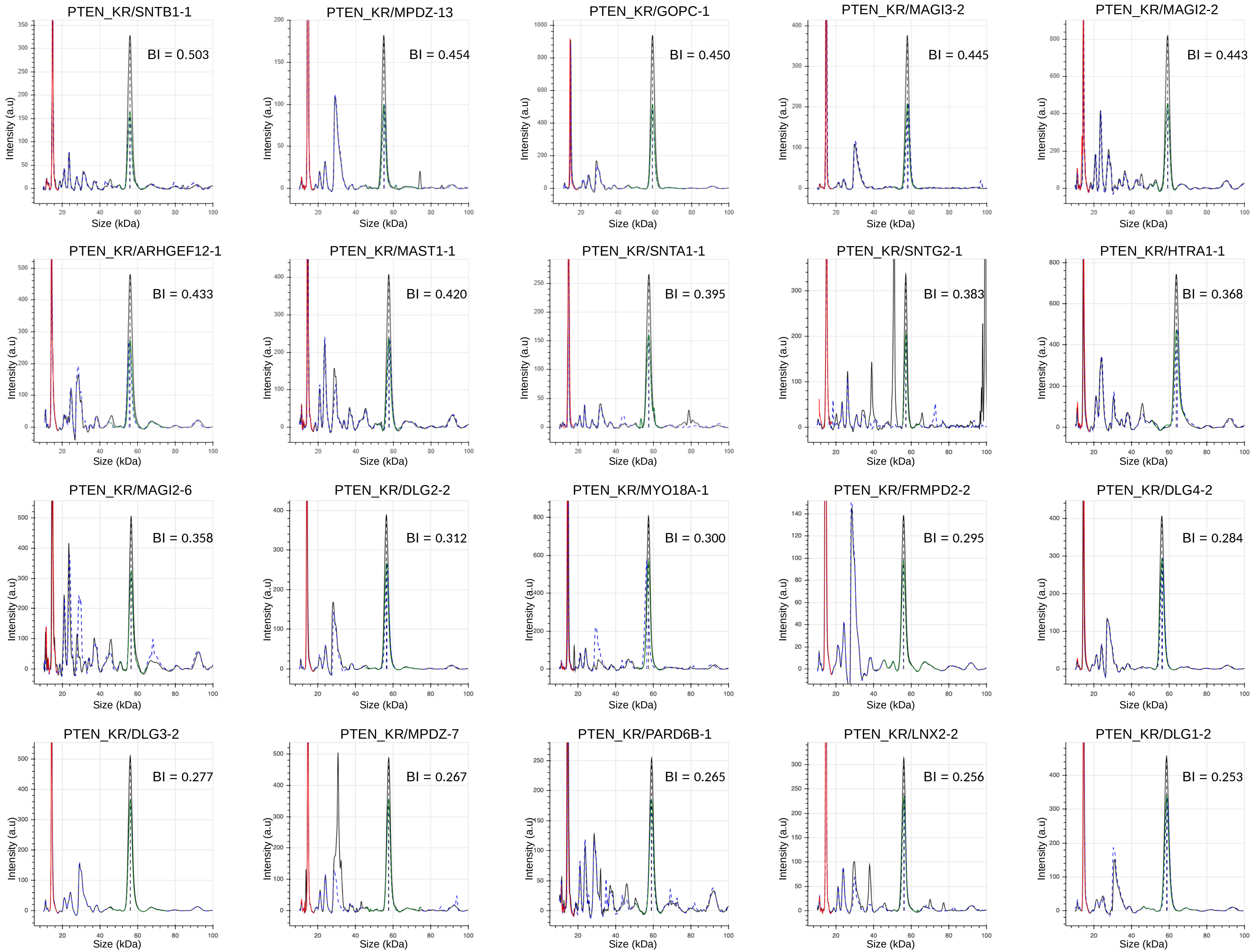

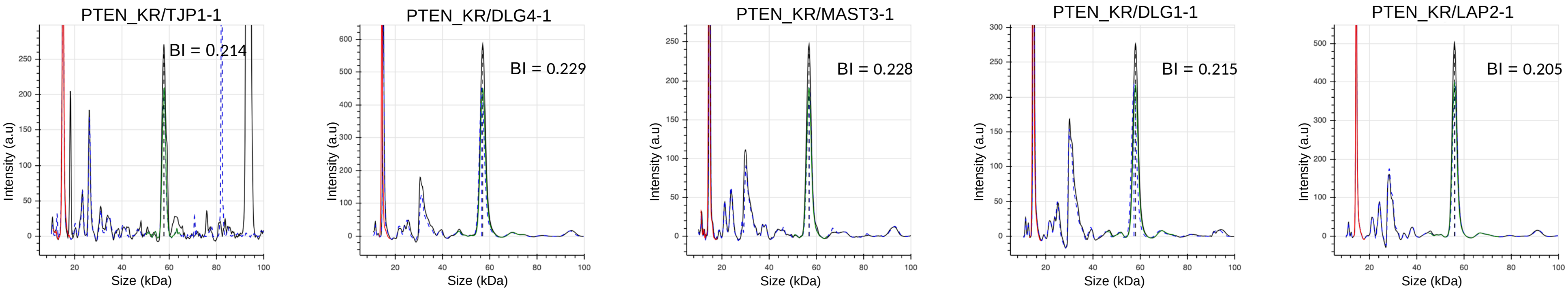

### Supp. Fig. S2

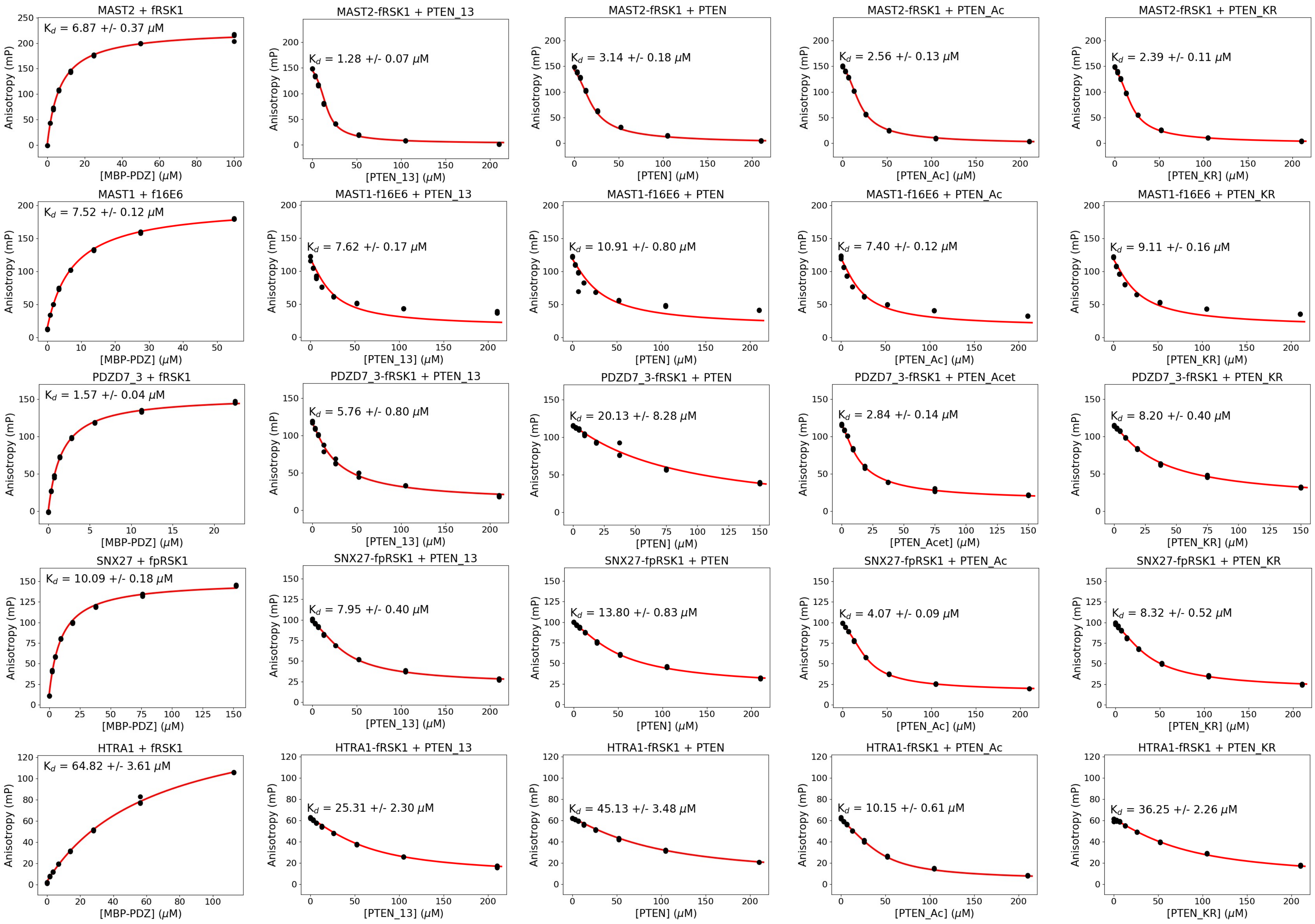

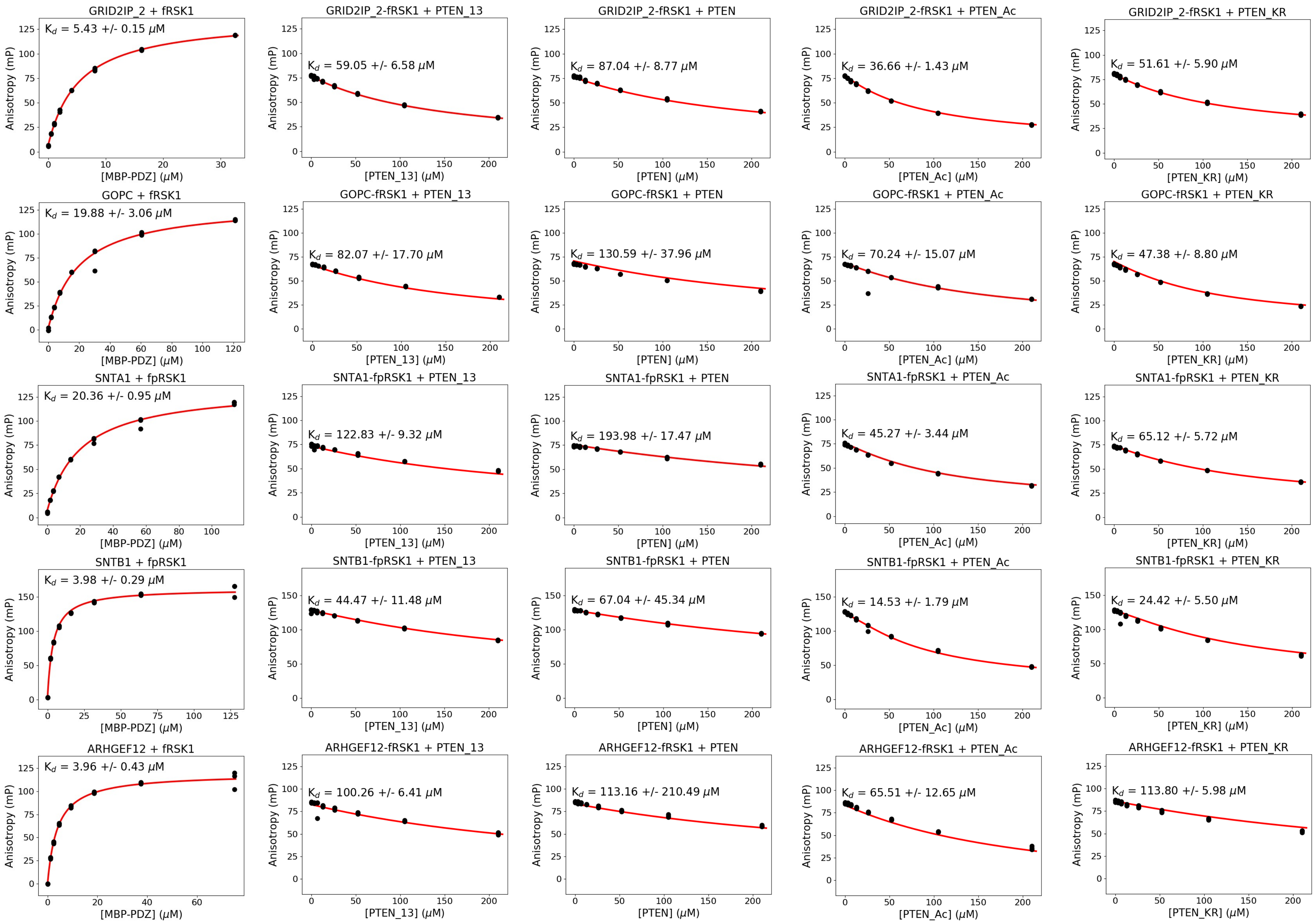

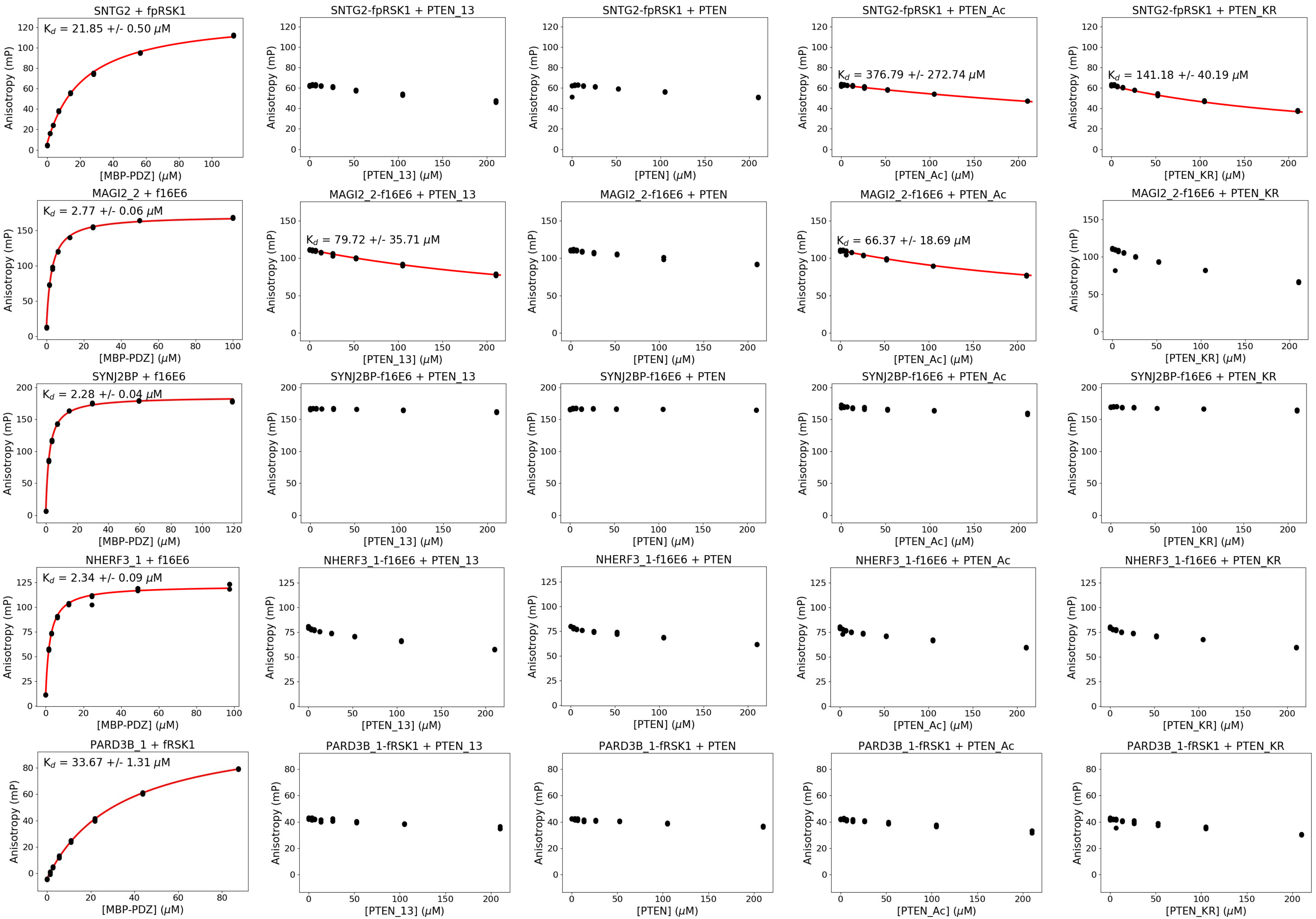

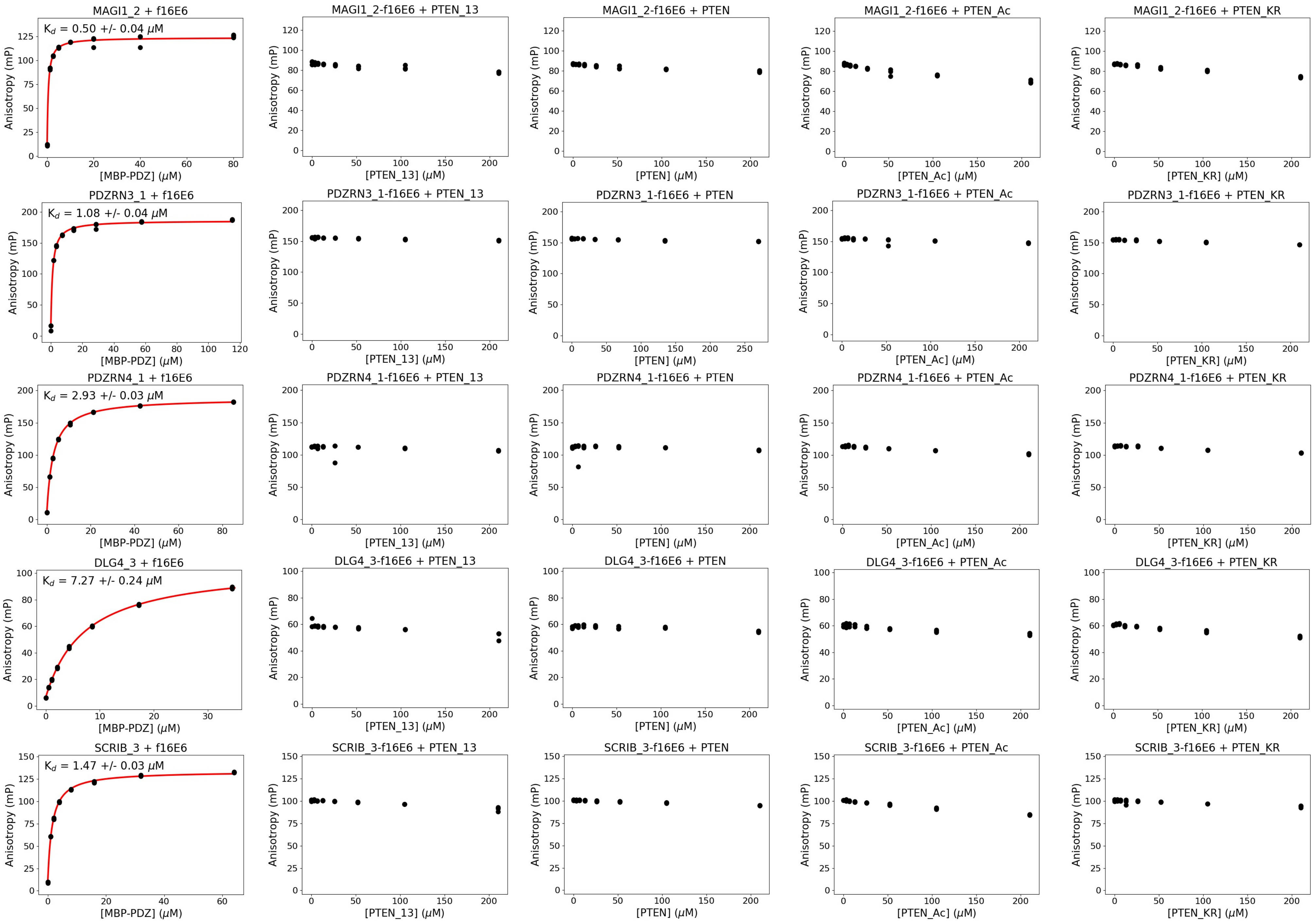

### Supp. Fig. S3

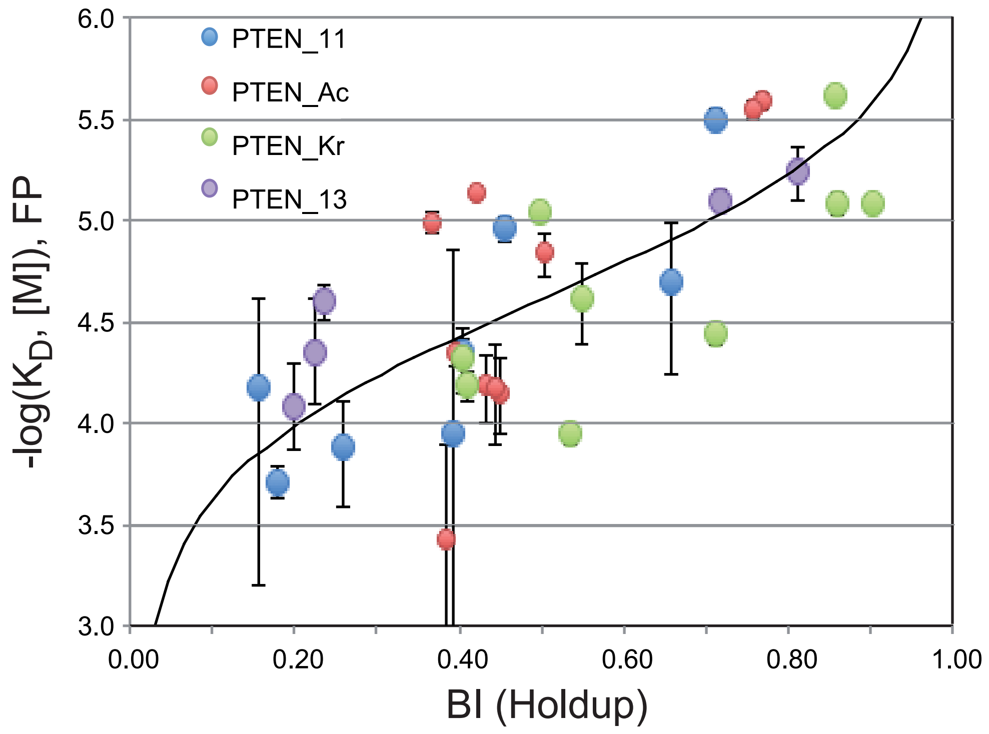
